# Neuronal Connectivity as a Determinant of Cell Types and Subtypes

**DOI:** 10.1101/2023.08.09.552547

**Authors:** Lijuan Liu, Zhixi Yun, Linus Manubens-Gil, Hanbo Chen, Feng Xiong, Hongwei Dong, Hongkui Zeng, Michael Hawrylycz, Giorgio A. Ascoli, Hanchuan Peng

**Author notes:** Correspondence: Hanchuan Peng.

## Abstract

Classifications of single neurons at brain-wide scale is a powerful way to characterize the structural and functional organization of a brain. We acquired and standardized a large morphology database of 20,158 mouse neurons, and generated a whole-brain scale potential connectivity map of single neurons based on their dendritic and axonal arbors. With such an anatomy-morphology-connectivity mapping, we defined neuron connectivity types and subtypes (both called “c-types” for simplicity) for neurons in 31 brain regions. We found that neuronal subtypes defined by connectivity in the same regions may share statistically higher correlation in their dendritic and axonal features than neurons having contrary connectivity patterns. Subtypes defined by connectivity show distinct separation with each other, which cannot be recapitulated by morphology features, population projections, transcriptomic, and electrophysiological data produced to date. Within this paradigm, we were able to characterize the diversity in secondary motor cortical neurons, and subtype connectivity patterns in thalamocortical pathways. Our finding underscores the importance of connectivity in characterizing the modularity of brain anatomy, as well as the cell types and their subtypes. These results highlight that c-types supplement conventionally recognized transcriptional cell types (t-types), electrophysiological cell types (e-types), and morphological cell types (m-types) as an important determinant of cell classes and their identities.

## Introduction

A mammalian brain is a complex network of tens of millions or more neurons and supporting cells that work together to carry out its functions (Purves, et al, 2019; Luo, 2015). These neurons form intricate modules within neuronal circuits and connectomes (Lichtman, et al, 2008; Sporns, 2011; Van Essen, et al, 2012; Seung, 2012). Cataloging brain-wide neuron types is recognized as a powerful way to understand the structural and functional organization of the brain (Peng, et al, 2021; Winnubst, et al, 2019). A definitive class of neuronal cells in a mammalian brain typically has a number of neurons that share multiple common attributes. Recent advances in classifying neurons usually rely on four major types of attributes: anatomical (Zeng and Sane, 2017; Muñoz-Castañeda, et al, 2021), physiological (Gouwens, et al, 2020), morphological (Peng, et al, 2021; Winnubst, et al, 2019), and molecular (Moffitt, et al, 2018; Zhang, et al, 202l; Zhang, et al, 2023), which are further complemented by other attributes such as lineage or developmental trajectory of these cells (Sebé-Pedrós, et al, 2018; Zhong, et al, 2020; Russ, et al, 2021).

Large-scale data acquisition and analyses at single-neuron resolution have succeeded in identifying neurons based on their 3-D registered soma-locations to the standard brain atlases. For mouse brains, the Allen Common Coordinate Framework (Dong, 2008; Wang, et al, 2020) and image-modality specific variations (Qu, et al, 2022) serve as standard atlases to index anatomical locations of neurons of interest. However, such soma-location cell types, or s-types, provide only an anatomical reference of the respective neurons, with marginal indication about the structural, physiological, molecular, and other attributes of neurons. Practically available techniques (Lee, et al, 2021; Lipovsek, et al, 2021) to record electrophysiological, morphological, and transcriptional properties of individual neurons have generated massive resources of these data modalities (Kalmbach, et al, 2021), enabling the analysis of electrophysiological cell types (e-types), morphological cell types (m-types), and transcriptional cell types (t-types) that further classify various s-types (Scala, et al, 2021).

The description of the morphology of neurons has been a crucial force to advance neuroscience since the time of Cajal (Cajal, 1909). While high-resolution digital reconstruction of the 3-D morphology of neurons is very challenging (Peng, et al, 2015; Manuben-Gil, et al, 2023), recent efforts in large-scale, semi-automatic reconstruction have yielded substantial datasets for whole mouse brains (Peng, et al, 2021; Winnubst, et al, 2019; Gao, et al, 2022) and other complex primate brains (e.g. Han, et al, 2023). We believe that a comparative approach taking advantage of these resources will help understand the morphological classification and distribution of single neurons. We also envision that an objective comparison of the morphology of neurons, especially from various data sources, should be carried out in a standardized coordinate system of an entire brain. Recent advances in brain mapping and registration (e.g., Qu, et al, 2022) provide such an opportunity.

Recent neuroscience research has highlighted the urgent need and various approaches to study neuronal connectivity and the whole-brain connectome (Abbott, et al, 2020; Whitesell, et al, 2021; Axer, et al, 2022). The MICrONS Consortium has recently produced a number of analyses about the connectivity of cortical neurons using electron microscopy (EM) datasets (e.g. Turner, et al, 2022; Dorkenwald, et al, 2022; Yin, et al, 2020). Parallel efforts also include the EM-based reconstruction and analysis of *Drosophila* hemibrain that also have a limited use of the cell connectivity to define cell types (Scheffer, et al, 2020). However, the EM approach has not yet scaled up to a whole mouse brain, which motivated us to take an alternative approach. Indeed, the connectivity of neurons has to be mediated by their morphology, making it challenging to study at the whole-brain scale. While it is clear that neurons can be classified based on their regional projection and connectivity, a quantitative study of the connectivity types of neurons in mammalian brains has been challenging. The goal of this study is to make an initial attempt to define neuronal connectivity in the context of whole brain based on aggregating and augmenting the largest single neuron morphology reconstruction datasets. In particular, we analyzed data with axonal and dendritic reconstructions of all brain regions to catalog connectivity types (or c-type) and subtypes of anatomically defined neurons. We further investigated the role of such connectivity types, and found that the connectivity types and subtypes supplement with the conventionally recognized transcriptional cell types (t-types), electrophysiological cell types (e-types), and morphological cell types (m-types), as a new determinant of cell classes and identities. Our finding underscores the importance of potential connectivity in characterizing the modularity of brain anatomy, as well as the cell types and their subtypes.

## Results

### Whole-Brain Map of Neuron Arbors and Projections

To formalize terminology, we call each group of neurons whose somas are in the same brain region a “soma-type”, or *s-type*. A neuron type determined based on clustering of morphological features, or *m-features*, is called a “morphology-type”, or *m-type*. Many morphological features, such as length, surface area, and number of branches of a neuron, are independent of a neuron’s spatial orientation, while other m-features, such as width and height, may be associated with a neuron’s orientation. A neuron type determined based on clustering of connectivity features/profiles, or *c-features*, of neurons is called a “connectivity-type”, or *c-type*. All *c-features* are orientation-independent and thus c-type is also not associated with a neuron’s orientation. Of note, connectivity is often associated with a topological direction, which means where axons project to and where neuronal input signal come from. Morphology features do not immediately exhibit such a directionality except partitioning into dendritic and axonal arbors. Remarkably, any quantifiable neuronal “connectivity” must be based on an anatomically precise mapping of individual neurons’ morphology at the whole brain scale. In another word, c-type is a derivative of m-types but also involves multiple neurons, neuron-populations, and brain regions, in a standardized manner.

To effectively study s-types, m-types, and c-types, we built a comparative morphology neuron database that consists of 20,158 neuron-morphology reconstructions (**Figure 1**), with which we inferred potential axon-dendrite connectivity of single neurons for standard whole-brain anatomical regions. To do so, we first aggregated four state-of-the-art neuron reconstructions datasets from independent sources (**Supplementary Table 1**). In detail, we specifically generated 3-D dendritic morphology of 10,860 neurons, called DEN-SEU (**Figure 1A** and **1B, Supplementary Figure 1**), to complement the full morphologies (complete axons and dendrites) of 1741 neurons in the BICCN AIBS/SEU-ALLEN neuron morphology dataset (Peng, et al, 2021) and 1200 neurons in the Janelia MouseLight project dataset (Winnubst, et al., 2019) (**Supplementary Figure 1**), and the axonal morphology of 6357 neurons generated by ION (Gao, et al, 2022). For fair and comprehensive analyses, we cross-validated neuron morphologies (**Supplementary Figures 2 and 3**) to avoid potential systematic bias favoring one particular way in generating the respective neuron data. Moreover, we applied 3-D brain registration (**Methods**) to all these neurons to map them onto the same spatial coordinate system, Allen mouse brain Common Coordinate Framework, CCFv3 (Wang, et al, 2020), so that all these neurons’ soma-locations and 3-D morphologies can be compared against each other directly.

**Fig 1.**
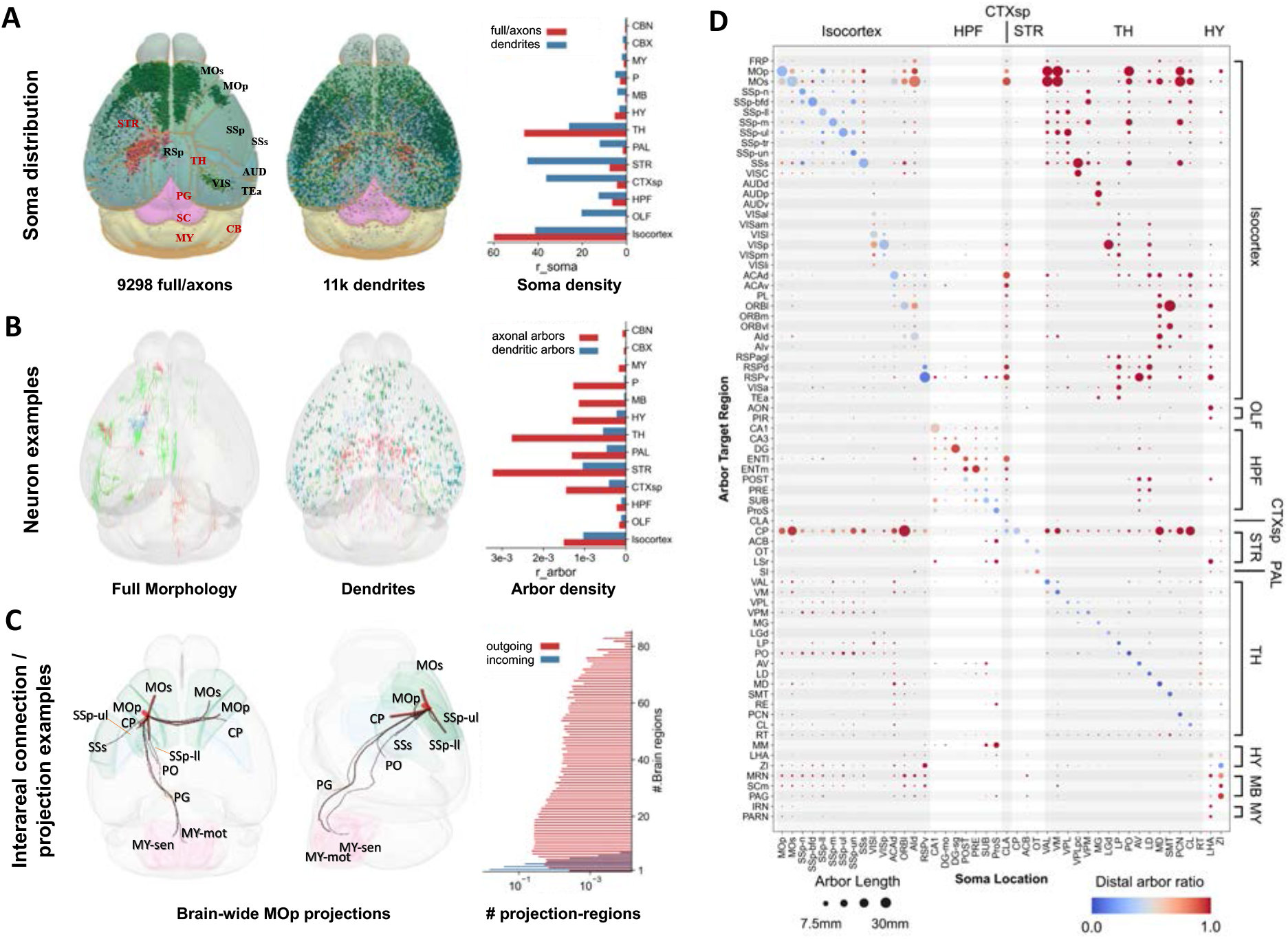
Overall distribution, arborization and potential connections of 3-D mapped and registered neurons at whole brain level. **A.** Standardized 3-D locations of 20,158 neurons in this study, pooled in two cohorts, i.e., one with full axon reconstructions (n=9298) and one with local dendritic reconstructions (n=10860) that covers all brain regions. Bar-chart: soma density (r_soma, per mm^3^) in main brain regions (see **Supplementary Table 2** for abbreviations). **B**. 38 full neuron reconstruction examples with different arborization patterns innervated from 8 brain regions (CTXsp, HPF, Isocortex, OLF, PAL, STR, TH, P, see **Supplementary Table 2** for abbreviations) and 20 dendritic reconstructions in different brain regions. Bar-chart: arbor density (r_arbor, per μm^3^) for major brain regions. **C.** Connection examples indicated by the projection patterns of primary motor cortex (MOp) cells to primary somatosensory cortex (SSp), SSp - lower limb (SSp-ll), SSp - upper limb (SSp-ul), secondary somatosensory cortex (SSs), caudate putamen (CP), secondary motor cortex (MOs), posterior complex - thalamus (PO), pontine gray (PG), medulla (MY), Medulla - sensory related (MY-sen), Medulla - motor related (MY-mot) regions. Bar-chart: histogram of outgoing and incoming connections of brain-wide projections; r_proj: the ratio of the number of neurons passing through a specific number of brain regions normalized against the total number of neurons. **D.** Whole brain arborization map of 2941 neurons. A similar map for all 9298 neurons with axons was also produced (**Supplementary Figure 6**). Horizontal axis: soma location of single cells. Vertical axis: arbor projection regions which are also grouped into larger brain areas. Size of circles: arbor length in brain regions. Color bar: ratio of local and distal arbors relative to soma locations.

The somas of the DEN-SEU neurons are distributed fairly evenly across major brain regions, while the other three full morphology or axon datasets focus on specific brain regions of cerebral cortex, thalamus, striatum, hypothalamus, hippocampus, and claustrum (**Figure 1A; Supplementary Table 2**). Compared with the sparse and long projection of axons in the other datasets, the dendritic arbors of DEN-SEU cover all CCFv3 brain regions (**Figure 1B**), making this dataset suitable for analyzing the target projection/connection regions of axons of neurons. We also cross-validated the quality and distribution of our assembled neuron data with other public documented neuron morphologies of shared by independent labs (**Supplementary Figures 4 and 5; Supplementary Table 3**) via NeuroMorpho.Org (Akram, et al, 2018; Bijari, et al, 2020). At the same time, the projection pathways of aggregated full neurons and axons data in this study capture many important regional connections, such as neurons originating from primary motor cortex (MOp) (**Figure 1C**). The total number of brain regions reached by projection axons follows a broad distribution (**Figure 1C**), indicating that most axons normally project to relatively distal regions. By contrast, dendrites extend a much shorter distance, invading at most 5 or 6 brain regions nearby their soma anatomical locations (**Figure 1C**). The large number of neurons involved in this study form complex patterns of potential connectivity, which should be quantified and analyzed in a principled way. We tackled this challenge by considering the modularity and granularity of individual neurons.

Neuron arbors often correlate with regions of dense connections between neurons. Therefore, we used a recent machine learning method, AutoArbor (Peng, et al, 2021), to determine the topologically connected arborization regions of neurons automatically. In particular, we started with 2941 fully reconstructed neuron morphologies in the Allen/SEU-ALLEN and MouseLight datasets to produce a brain-wide arborization map of a mouse brain (**Figure 1D**). In this way, various neuronal pathways indicated in our datasets (**Figure 1C**) are quantitatively modeled. For example, we observed clear modules of projection and potential connection patterns in large brain regions such as isocortex, striatum, and thalamus. This motivated us to characterize the potential connectivity among neurons using the structural components, i.e., neurite arbors, systematically. Accordingly, we generated in total 26205 axonal arbors and 20158 dendritic arbors (**Figure 1B**) for all neurons in this study. We subsequently used these arbors to define the connectivity among neurons and respective c-types.

### Connectivity Profiles Augment Morphology-Based Neuron Types

Distinct from morphology analyses of neurons that rely on various m-features, such as the Sholl analysis (Binley, et al, 2014), L-Measure (Scorcioni, et al, 2008), and extended global or local structural features (Wan, et al, 2015), we study cell-typing by generating the neuron connectivity features. One approach to quantifying single neuron connectivity is based on axon-dendrite colocalization that needs precise details on synaptic contact location approximation (Rees, et al. 2017), which however is still challenging at the whole brain scale. Our intuitive approach is to use soma locations and defined spatial domains of neuronal arborization. Specifically, we determined the connection targets of a neuron based on the 3-D registered brain regions invaded by its axonal arbor, and the connection strength based on the spatial adjacency of this neuron’s axonal arbor and nearby dendritic arbors of neurons in our dataset (Figure 2A). We detected arbor domains of neurons that originate from a specific brain region using Gaussian mixture models (**Methods**) and produced spatially and statistically optimal parcellation of projection sites of all s-types. Within each arbor domain, the arborization pattern of each group of neurons of the same s-type is approximated using a spatially homogeneous Gaussian distribution. For example, SSp neurons were found to have 9 arbor domains, 4 of which are axonal arbor domains and 5 are dendritic arbor domains (Figure 2B).

**Fig 2.**
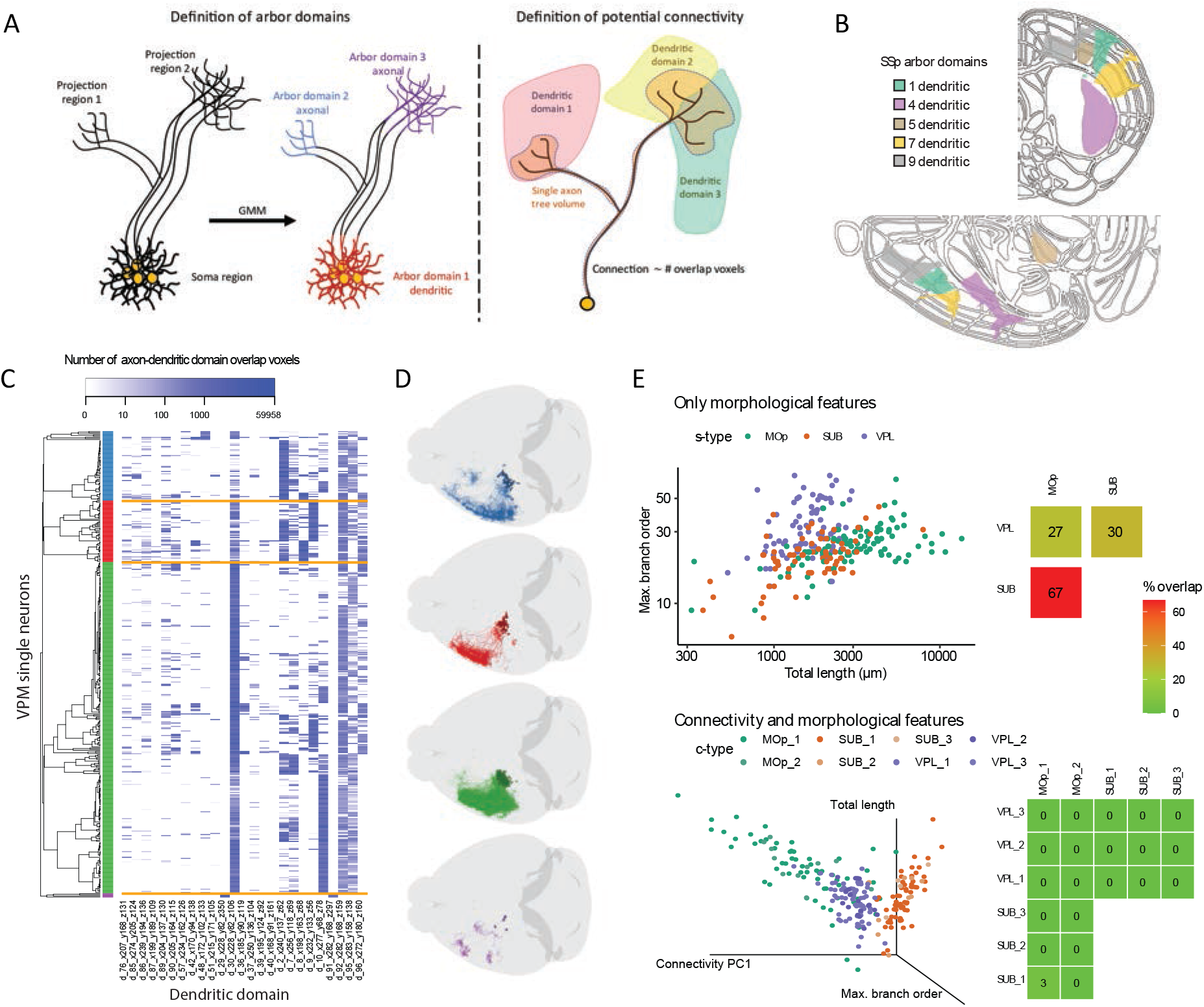
Formulation of neuronal connectivity and difference between morphological features and connectivity features of single neurons. **A.** Schematic overview of the definition of arbor domains and potential connectivity. (Left) Neurons belong to the same soma-location type (s-type) when their cell bodies are located in the same Allen brain atlas Common Coordinate Framework (CCFv3) anatomical region. In each s-type, the neuron morphological coordinates are spatially clustered using a Gaussian Mixture Model (GMM). Each resultant cluster forms an arbor domain. A dendritic arbor domain contains a major number of somas. Axonal arbor domains: any non-dendritic domains. (Right) Overlapping voxels between axonal and dendritic domains define the potential connectivity. **B.** Exemplar arbor domains for SSp neurons in middle sections of the CCFv3 atlas outline (top, coronal half-view; bottom, sagittal half-view). Note that the axonal arbor domains are not shown for clarity. **C.** Heatmap of potential connectivity for VPM (ventral posteromedial nucleus - thalamus) neurons, which project to SSp heavily. Horizontal axis: dendritic domains (as indicated by the prefix ‘d’) with renumbered identifier denoting the domain center coordinates in **B** (see **Supplementary Table 4** for a complete list of domains); only the top-25 domains with the greatest variances are shown for clarity while the entire feature vector was used in clustering. Vertical axis: clustered VPM neurons. Dendrogram in the left: hierarchical clustering (four clusters in blue, red, green, and purple) of the potential connectivity feature vectors of neurons. Orange lines: cluster boundaries in the heatmap. Color bar: the number of overlapping voxels between a neuron-of-interest and dendritic domains. **D.** Horizontal view of VPM neurons overlayed on the CCFv3 contour colored by the clusters obtained from potential connectivity. **E.** Comparison of clustering results based on morphology features only (top) and based on joint feature vectors by concatenating morphology and connectivity features (bottom). (Top left) Scatterplot of MOp (green), SUB (subiculum; dark orange) and VPL (ventral posterolateral nucleus - thalamus; purple) s-types. Horizontal axis: total length of the neurons in μm. Vertical axis: maximum branch order. (Bottom left) 3-D scatter plot of the total length, maximum branch order, and the first component of a Principal Component Analysis (PCA) of the potential connectivity matrix. The c-types obtained are colored with different shades of each s-type color (e.g., MOp_1 dark green, MOp_2 light green). (Top and bottom right) Heatmaps of the overlap between point clouds in the scatterplots. Color bar: percentage of overlap between s-type pairs, measured by misclassified neurons when using a Support Vector Machine (SVM) to classify the data.

For each neuron in a specific s-type, we then computed a connection barcode (Figure 2C) as the features to characterize the axon-dendritic spatial overlap of its axonal arbors and dendritic arbor domains of all s-types at the whole brain scale, all defined in the standard CCF space. For DEN-SEU, we produced 19 dendritic arbor domains per brain hemisphere. We also produced another 56 dendritic arbor-domains per brain hemisphere for other s-types with at least 60 reconstructed neurons. These dendritic arbor domains span an average volume of 8.94 mm^3^. The resultant connection barcode is thus a 150-dimensional feature vector for the entire brain, indicating how axons of neurons in a s-type may project and potentially connect to various dendritic domains in the context of whole brain anatomy. With this barcode, neurons belonging to a s-type can be further clustered. For instance, ventral posteromedial nucleus (VPM) neurons were smoothly clustered into four connection groups (Figure 2C), which are visually separable from each other (Figure 2D).

To understand the advantage of the connectivity barcode, we first applied it to assisting conventional morpho-analysis of cell types that clusters s-types or their sub-types based on m-features. It was difficult to separate MOp, Subiculum (SUB), and ventral posterolateral nucleus (VPL) neurons that have heavily overlapping m-features, as seen in both the overlap scores and the feature scatter plot (Figure 2E top). However, when the connectivity features were appended to the m-feature vectors to cluster these three s-types, they became clearly separable in terms of a minimal overlapping in this case (Figure 2E bottom right). When the first principal component of the connectivity features was added in visualization, the separation of the three s-types was visible (Figure 2E bottom left). This shows that the connectivity features help discriminate neuron classes, similar to the dimension-increment analysis or support vector machines (Cortes and Vapnik, 1995; Steinwart and Christmann, 2008) in pattern recognition and machine learning, where non-separable classes could become distinguishable in higher-dimensional spaces.

### Connectivity Types Outperform Morphology Types in Neuron Classification

To investigate whether c-features would classify cell types better than conventionally used m-features (Zeng and Sane, et al, 2020; Peng, et al, 2021), instead of providing auxiliary dimensions to assist cell typing, we computed morphological features’ similarity scores (m-score) of all 31 known s-types (*n* > 60) (Figure 3A). Except a small amount, i.e., 25.8%, of s-types that have relatively low similarity in their m-features, the majority of s-types (74.2%) make up 3 boxed cohorts, within each of which neurons of different s-types share high similarity m-features (Figure 3A). Remarkably, the similarity score between the c-features (c-score) of all 3 cohorts of s-types are dramatically reduced while the c-scores of the other 8 s-types remain low (Figure 3B). In another word, in general a s-type is well separated from other s-types in the space of c-features.

**Fig 3.**
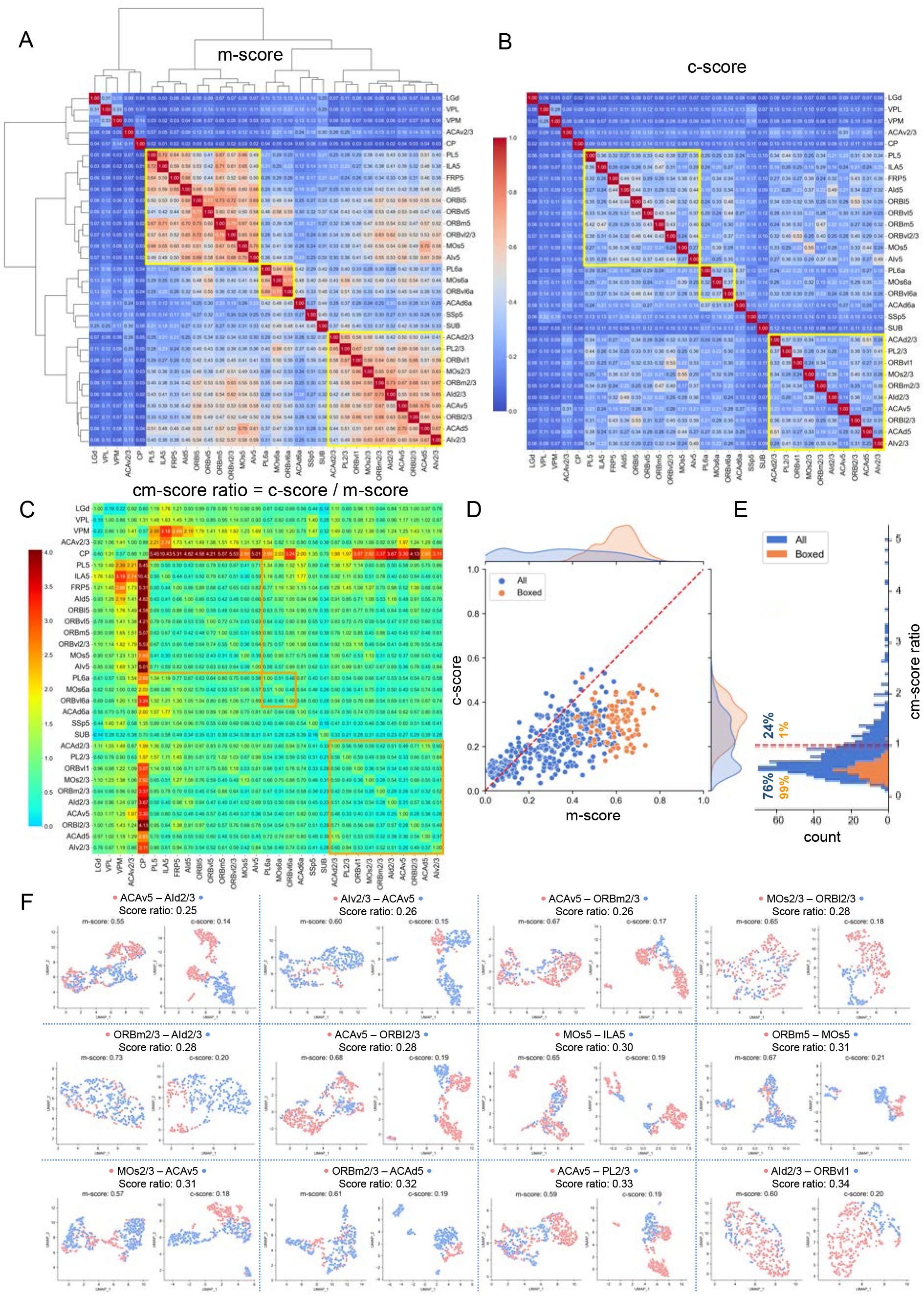
Classification of cell types based on morphological and connectivity features. **A.** Clustering based on similarity score of morphology features, i.e., m-score, of 31 s-types (n>60 in each) in cortex, thalamus, and striatum. Yellow boxes: 3 cohorts of s-types that share highly similar morphology features. Values in matrix: normalized similarity between 0 and 1. **B**. Similarity score of connectivity features, i.e., c-scores, sorted using the same order of s-types as in **A**. Color bar: normalized similarity among features (also the same as in **A**). **C.** Ratio matrix of c-score in **B** over m-score in **A**. **D**. Joint and marginal distributions of corresponding c-scores and m-scores for all pairs s-types (blue) and boxed pairs in **A** and **B** (yellow). **E**. Histogram of c/m-score ratios in **C**. **F.** Paired comparison of UMAP clustering of s-types using either morphology or connectivity features, corresponding to the 12 smallest c/m-score ratios in **C**.

We further directly compared corresponding m-scores and c-scores to quantify the improvement of cell typing performance of connectivity features over morphological features. Here, 76% of entries in the ratio matrix of c-scores and m-scores (Figure 3C**, 3D, and 3E**) are less than 1, while 99% of such entries corresponding to the boxed cohorts are less than 1. We also visualized the actual clustering of neurons based on either morphological features or connectivity features. Examination of the paired UMAP (Uniform Manifold Approximation and Projection) clustering for the 12 smallest ratios of c-scores and m-scores shows that c-features are much more separable than m-features (Figure 3F). For example, ACAv5 neurons have mixed m-features with AId2/3 neurons, however their c-features are clearly separable (Figure 3F). This is the same case for MOs5 neurons vs ILA5 neurons, ORBm2/3 neurons vs ACAv5 neurons, and all other visualized pairs of cell types, although all these cases have varying distributions in their UMAP space (Figure 3F). As our results analyze the largest neuron archives for the mouse brain containing major neurons classes, it is reasonable to conclude that c-features could serve as strong contenders of m-features for cell typing of neurons whose somas are from well-established anatomical regions.

### Connectivity Features Correlate with Spatial Separation of Potential Cell Subtypes

After establishing that connectivity is a powerful attribute for classifying neurons types, we investigated whether c-features would also help identify sub-types of neurons that share their soma locations in the same anatomical area. To do so, we generated a distance map (d-map) to measure the spatial separation of two neurons based on their soma locations (Figure 4A). Because within any specific brain region neurons were labeled in a stochastic way, the pairwise soma-distance may form a Gaussian-like or Gaussian-mixture distribution (Figure 4B). Particularly, when somas scatter almost uniformly within a brain region, their pairwise distance will be close to Gaussian, such as LGd and CP neurons (Figure 4B, red and blue). Conversely, when somas form two or more subclusters within a region, their pairwise distances may form a distribution with long-tail, or approximately a Gaussian mixture distribution, such as the ACAd 6a neurons in this database (Figure 4B, green). Correlating the morphology similarity scores (m-scores) and connectivity similarity scores (c-scores) with d-map provides a useful way to understand which kind of features may help identify subtypes of neurons whose somas are from subareas in an established s-type.

**Fig 4.**
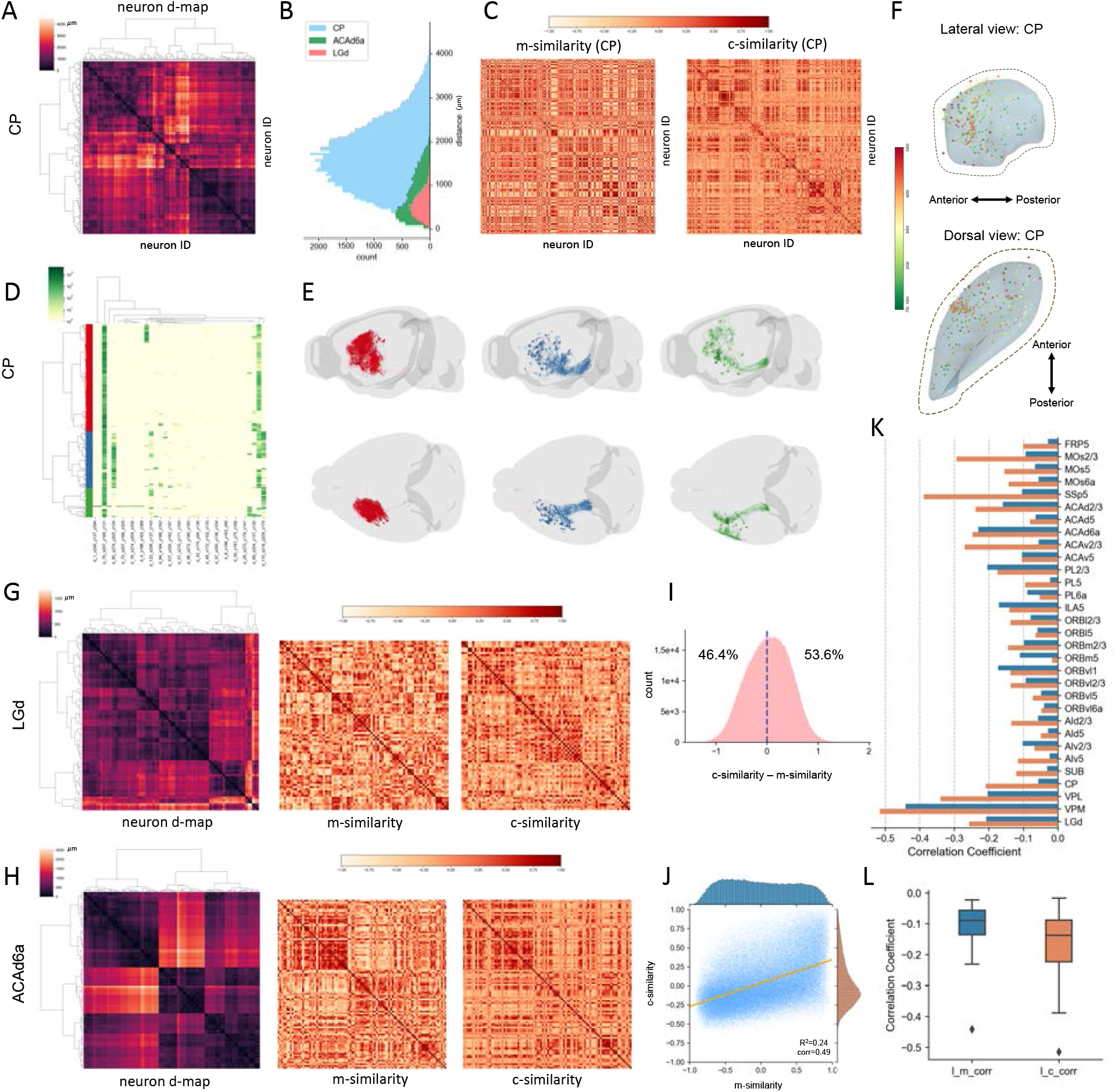
Classification of cell subtypes based on morphological and connectivity features. **A.** Pairwise soma-distance map of CP neurons bi-clustered based on spatial adjacency of somas mapped to CCFv3. **B.** Histograms of the pairwise soma-distances for neurons in CP, ACAd6a (Anterior cingulate area, dorsal part, layer 6a) and LGd (lateral geniculate complex - dorsal part) regions. C. Matrices of morphology-feature similarity scores (m-similarity) and connectivity-feature similarity scores (c-similarity) of individual CP neurons, rows and columns sorted in the same order of the clustered distance map in **A**. Cosine similarity scores are used. **D.** Connectivity-feature based clustering of CP neurons into two main subclasses (red and blue), in the same convention as Fig. 2C. **E**. 3-D visualization of the two CP neuron-subclasses in **D**. **F**. 3-D soma-locations of CP neurons. Color: the largest Euclidean distance between axonal terminals and the respective soma. **G**. LGd neurons’ distance map and respective m-and c-similarity matrices, rows and columns sorted in the same order. **H**. ACAd6a neurons’ distance map and respective m-and c-similarity matrices, rows and columns sorted in the same order. **I**. Histogram of the difference between corresponding c-and m-similarities for all neurons of the 31 s-types in this study. **J**. Scatter plot and marginal distributions of corresponding c-and m-similarities for all neurons in the 31 s-types. **K**. Correlations between soma-distance-map and c-or m-similarity for all 31 s-types. **L**. Overall correlations between neurons’ soma distances and the respective similarities in connectivity features or morphology features. l_m_corr: correlation between location-distances and m-similarities. l_c_corr: correlation between location-distances and c-similarities.

In the example of CP neurons, we calculated the pairwise m-score and c-score matrices (Figure 4C) sorted in the same order of neurons as in the respective d-map (Figure 4A). Using the c-features, we obtained three major CP clusters (Figure 4D) with different projection and arborization patterns (Figure 4E), although their somas are mixed fairly uniformly (Figure 4F), while there is no obvious subcluster based on m-features’ similarity matrix (Figure 4C). Similarly, we computed the d-maps and respective m-scores and c-scores matrices for LGd and ACAd6a neurons (Figure 4G, Figure 4H). The Gaussian-mixture like distribution of the pairwise neuron-distances of ACAd6a neurons also translate to potential clusters in ACAd6a’s d-map (Figure 4H), while the single Gaussian-like distributions of CP and LGd neurons (Figure 4B) correspond to the less clear hierarchical clustering of the respective sorted d-maps (Figure 4A, Figure 4G).

We computed the corresponding d-maps, m-score, and c-score matrices for all 31 s-types of neurons. Overall, we found that for any pair of neurons, their c-scores are only slightly greater than the m-scores (Figure 4I). There is a weak positive correlation between these two scores (Figure 4J), of which m-scores follow a much flatter marginal distribution than c-scores (Figure 4J); this indicates that statistically it would be harder to produce clearly segregated neuron clusters based on morphology similarity. However, remarkably the corresponding entries of the d-map and c-score matrices have evidently negative correlation, which is also much stronger than that between d-map and m-score entries (Figure 4K, Figure 4L). Indeed only 6 out of 31, or 19.4%, s-types show stronger negative-correlation of soma-location-and-morphology similarity over soma-location-and-connectivity correlation (Figure 4K). Neurons with far away soma locations can be at most 4 times more likely to have different c-features than m-features (Figure 4L). Thus, we conclude that potential subtypes for a s-type are statistically better represented by c-features than by m-features.

### Spatially Tuned Connectivity Features Identify Cell Subtypes

Anatomical sub-grouping of neurons within a specific brain region reflects the spatial coherence of these cells. As c-features correlate more strongly with the spatial adjacency of neurons, for each s-type we combined connectivity profiles and spatial adjacency to cluster neurons and identify potential anatomical subtypes. We called this approach Spatially-Tuned c-Features, with which we produced clear subtyping of neurons (Figure 5) that we had never been able to identify using alternative methods.

**Fig 5.**
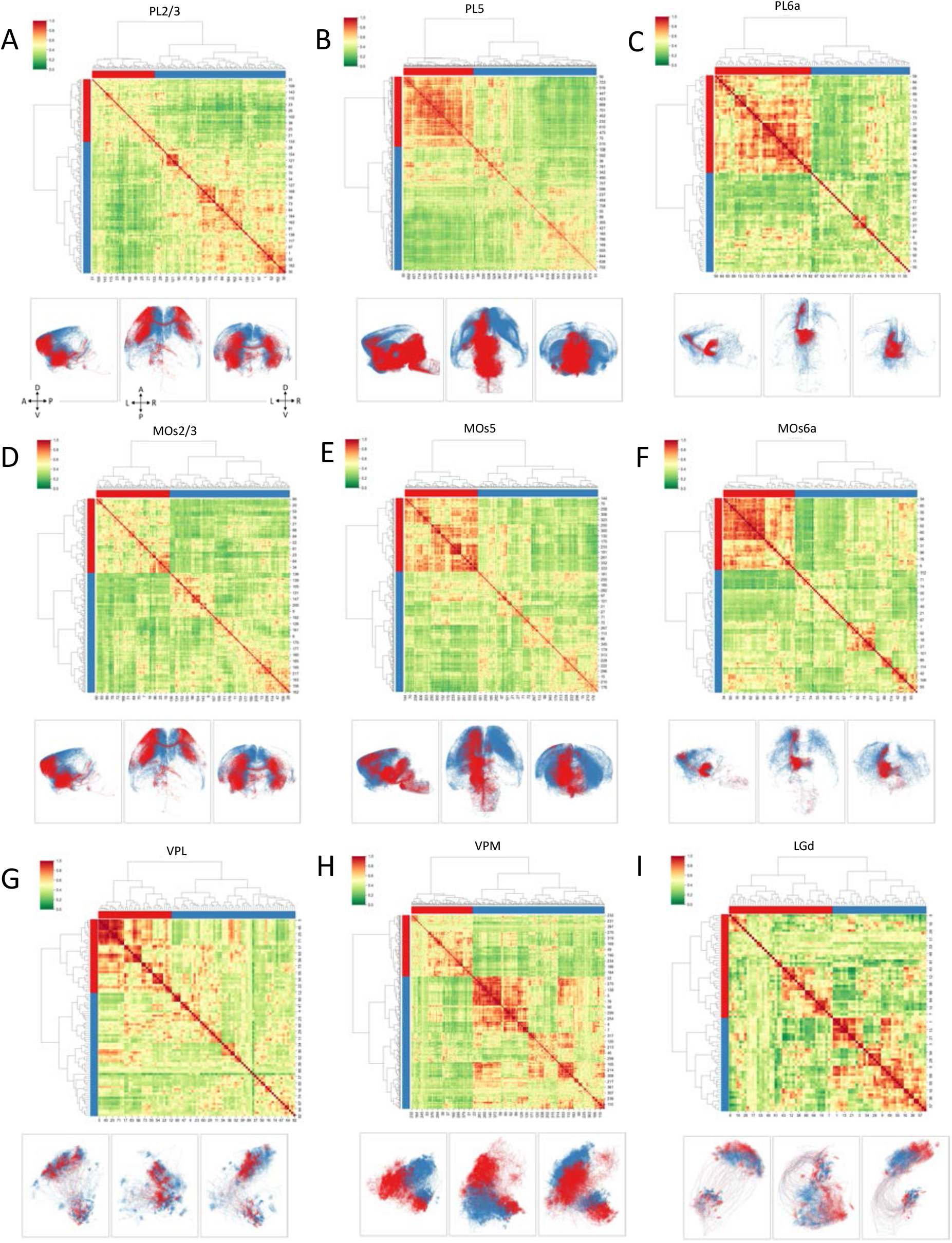
Neuron subtyping based on spatial-connectivity patterns. In each image, Upper row: bi-clustered spatially tuned connectivity similarity matrices, where different colors along the x-and y-axes indicate the clusters, and the index-numbers of neurons in a specific s-type are shown in both x-and y-axes. Lower row: tri-view visualization of neurons in CCFv3; neurons are rendered in the same colors as in the respective upper-row clusters. A: Anterior, P: Posterior, D: Dorsal, V: Ventral, L: Left, R: Right. **A. B.** and **C.** Subtyping of PL (prelimbic area -prefrontal cortex) neurons for layer 2/3 (PL2/3) (n=188), layer 5 (PL5) (n=795), and layer 6a (PL6a) (n=99), respectively. **D. E.** and **F.** Subtyping of secondary motor cortex neurons in MOs layer 2/3 (MOs2/3) (n=218), layer 5 (MOs5) (n=359), and layer 6a (MOs6a) (n=116), respectively. **G. H.** and **I.** Subtyping of thalamic neurons in VPL (n=91), VPM (n=406), and LGd (n=78), respectively.

In particular, for cortical neurons (Figure 5A**∼F**), we found that neurons in the prelimbic area (PL) have 2 subtypes for each of the layers 2/3 (Figure 5A), layer 5 (Figure 5B), and layer 6a (Figure 5C), respectively. Neurons in layers of the secondary motor cortex, MOs, could also be clustered into subgroups (Figure 5D**∼F**). The layer 2/3 MOs neurons are clustered into two large subgroups indicated by the sorted distance matrix, along with distinct projection patterns of these subgroups in the cross-sectional views of the CCF space (Figure 5D). Similarly, each of layer 5 and layer 6 MOs neurons were divided into two subgroups, respectively (Figure 5E **and 5F**). Detailed examination of these MOs subtypes provides guidance for analyzing connectivity-based subtypes of cortical neurons (see next section).

We also attempted to identify subregions in the thalamic gateway related to sensory and motor input, particularly VPL (Figure 5G), VPM (Figure 5H), and LGd (Figure 5I). We found that subregions of somas in these areas correspond to neurons projecting to distinguishable spatial targets, visualized often as homogeneous color-blobs of neuron-subclusters, which are particularly clear in the three subtypes of VPM neurons (Figure 5G). LGd has three known anatomical subregions (Guido, et al, 2018; Okigawa, et al, 2021), i.e., LGd-shell, LGd-core, and LGd-ip (ipsilateral zone). We found two major distinguishable subtypes of somas using our approach, which may provide further spatial granularity to study the previously documented subregions. Of note, LGd neurons could not be clearly clustered using either morphology features or connectivity or spatial distance features alone (Figure 4G).

### Subtyping MOs and VISp Neurons Reveals Diversified Connectivity, Transcriptomic, and Electrophysiological Characteristics

MOs neurons have long axonal projections that subserve animal decisions (Yang and Kwan, 2021). In addition to individual neurons’ spatial patterning, we also profiled the symmetry of MOs connectivity using the cortical layer 5 neurons. To do so, we kept the somas separated when calculating their space distance map. Our examination of individual MOs neurons confirmed long-range projection targets at the full-brain scale (Figure 6A, top). The overall projection patterns of these MOs neurons are also consistent with the previously documented population projection (Oh, et al, 2014) (Figure 6A, bottom-right). We found that the somas in MOs5_1 and those of MOs5_2 and MOs5_3 clusters distribute on the two sides of the brain (Figure 6A, bottom-left), while the somas in MOs5_2 and MOs5_3 essentially intermingled. Indeed, the projection patterns of MOs5_1 match well with the mirrored sum pattern of MOs5_2 and MOs5_3. In another words, the spatially tuned connectivity analysis provides a powerful way to reveal both the anatomical distribution of neuron subtypes and their symmetry. Particularly, while the reconstructions of neurons of this MOs5 dataset have three anatomical subtypes when both hemispheres of the brain are considered, there are only two genuine subtypes (Figure 5E) that are distributed symmetrically on the brain’s coronal plane. These two subtypes might be further subdividable as implied in respective clustering tree (Figure 5E).

**Fig 6.**
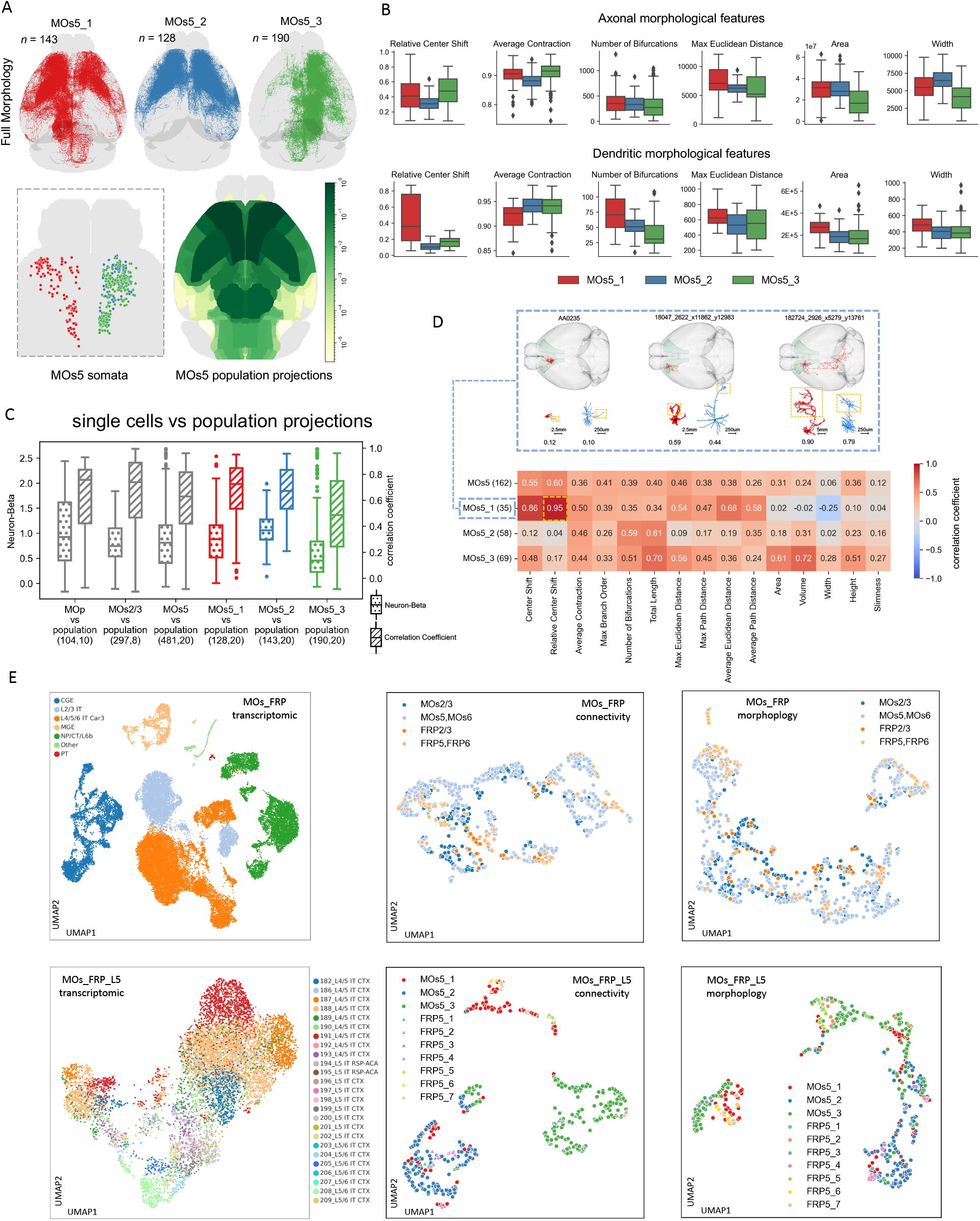
Comparison of various cell typing methods for MOs neurons. **A.** Three connectivity-based clusters for MOs5 neurons (top row) along with the distribution of their somas (bottom-left) and the overall projection patterns of MOs5 neurons (bottom-right) (Harris, et al, 2019). **B.** Key morphological features of the three connectivity-based MOs5 subtypes. **C.** Two metrics, neuron-beta and correlation coefficient, between single neurons and neuron-populations in motor cortex, specifically MOp, MOs2/3, MOs5, and MOs6a subtypes. **D.** Correlation of dendritic and axonal morphological features for MOs5 connectivity subtypes, along with examples of the first MOs5 cluster. Note that the clustered neurons in **A** might not have dendrite reconstructions, however in this dendro-axonal correlation analysis only neurons in **A** but also with full dendrites and axons are counted. **E.** Transcriptomic profile-based single neuron clustering of FRP-MOs neurons (n=34,331) and more specific FRP-MOs layer 5 neurons (n=9879), compared with the clustering based on connectivity and morphology features of FRP-MOs / FRP-MOs layer 5 neurons.

We also examined both the axonal and dendritic morphologies of MOs5 subtypes. While the most dendritic features of the two genuine subtypes, MOs5_2 and MOs5_3, are similar to each other, their axonal features (Figure 5E) are clearly different in area, width, and relative shift of centers, despite the similar numbers of axonal bifurcations. This means that, although these two subtypes have similar branching complexity, their projection patterns differ. Such variability of MOs5_2 and MOs5_3 is also seen in the different correlation and neuron-beta (Peng, et al, 2021; **Methods**) values compared to the overall MOs population projection (Figure 6C). The respective scores of MOs5-vs-population and MOs5_1-vs-population are comparable to each other, indicating MOs5_1 is a good ipsilateral approximation of the overall MOs5 patterns, also as the “sum” reference for MOs5_2 and MOs5_3 (Figure 6C). The MOs5 and MOs2/3 neurons also covary strongly with the MOs population projection. Differently, MOp neurons show more variation in the single-neuron-vs-population comparison, while their integrative projection pattern also matches with previous population projection data (Oh, et al, 2014) (Figure 6C). We also correlated the m-features of individual neurons’ dendritic and axonal arbors. For MOs5 neurons, m-features such as the number of bifurcations and total length show recognizable level of correlation, in the range of 0.3∼0.7, between dendrites and axons (Figure 6D).

We also produced UMAP analyses to compare the transcriptomic subtypes of single MOs neurons (Yao, et al, 2021), connectivity subtypes and morphological subtypes (Figure 6E). As the transcriptomic data of MOs and FRP (frontal pole, cerebral cortex) were mixed due to the limited spatial resolution at this point, we prepared connectivity and morphological features of individual neurons in a similar way, also specifically for layer 5. Within each of the individual scenarios, we observed relatively coherent subtyping except for the cases of morphological features. However, it seems that the diversity exhibited in the c-features cannot be immediately explained by the subgrouping of the transcriptomic features. We have not observed a conclusive layer-by-layer correspondence between transcriptomic and connectivity subtypes, either.

Moreover, we performed a joint analysis of the m-type, c-type, t-type and e-type data based on retrieving the publicly available electrophysiological and transcriptomic recordings of single neurons that also fall into the brain regions used in this study. For the primary visual area (VISp), we analyzed 48 fully reconstructed neuron morphologies and their regional connectivity patterns (**Supplementary Figure 7A**), along with their morphometric features (**Supplementary Figure 7B**) and anatomical locations of cell bodies (**Supplementary Figure 7C**). We found that VISp neurons in different cortical layers have a less clear separation in morphology (Supplementary Figure 6D) than in connectivity (**Supplementary Figure 7E**). We also re-analyzed previous single neuron electrophysiological recordings (Gouwens, et al, 2020) based on the concatenated e-type features (**Supplementary Table 7**) and colored these e-type data using their molecular profiles and anatomical locations (**Supplementary Figure 7F-G**). While it is clear that certain neurons in different layers have preferences in their physiological and molecular properties, there is a general disparity between such features and their c-types.

### Subtyping Single-Cell Connectivity of VP Nuclei Indicates Broader Multisensory Integration

By subtyping single cell reconstructions of 390 VPM and 83 VPL neurons, we were able to document the broad regional connections of VP neurons in a comprehensive manner. First, we clustered individual neurons’ detailed projections onto cortical areas and layers into 8 subtypes as a matrix (Figure 7A). These 8 groups have similar separation of their soma locations as well as the respective axonal arbor targets’ locations (Figure 7B **and 7C**). The longest dendrite can be about 5 times of the shortest dendrites in these groups (Figure 7D). We also confirmed the majority projection of VP neurons to layer 2/3 and 4 of somatosensory cortex (Figure 7A) consistent with previous knowledge at the neuron population level (e.g., Bureau, et al, 2006; Viaene, et al, 2011; Clascá, et al, 2012; Staiger and Petersen, 2021). It is interesting to note that, while our previous study (Peng, et al, 2021) implied that a small portion of VP projection may target MOp, the detailed examination presented in the next section (Figure 8) visualizes abundant outgoing arborization of VP neurons in MOp regions. Strikingly, we estimated a non-negligible 20.7% VP cells (*n* = 98) actually project to multiple cortical areas such as motor or visceral areas that are outside somatosensory cortex, and even beyond such as CP (Figure 7A).

**Fig 7.**
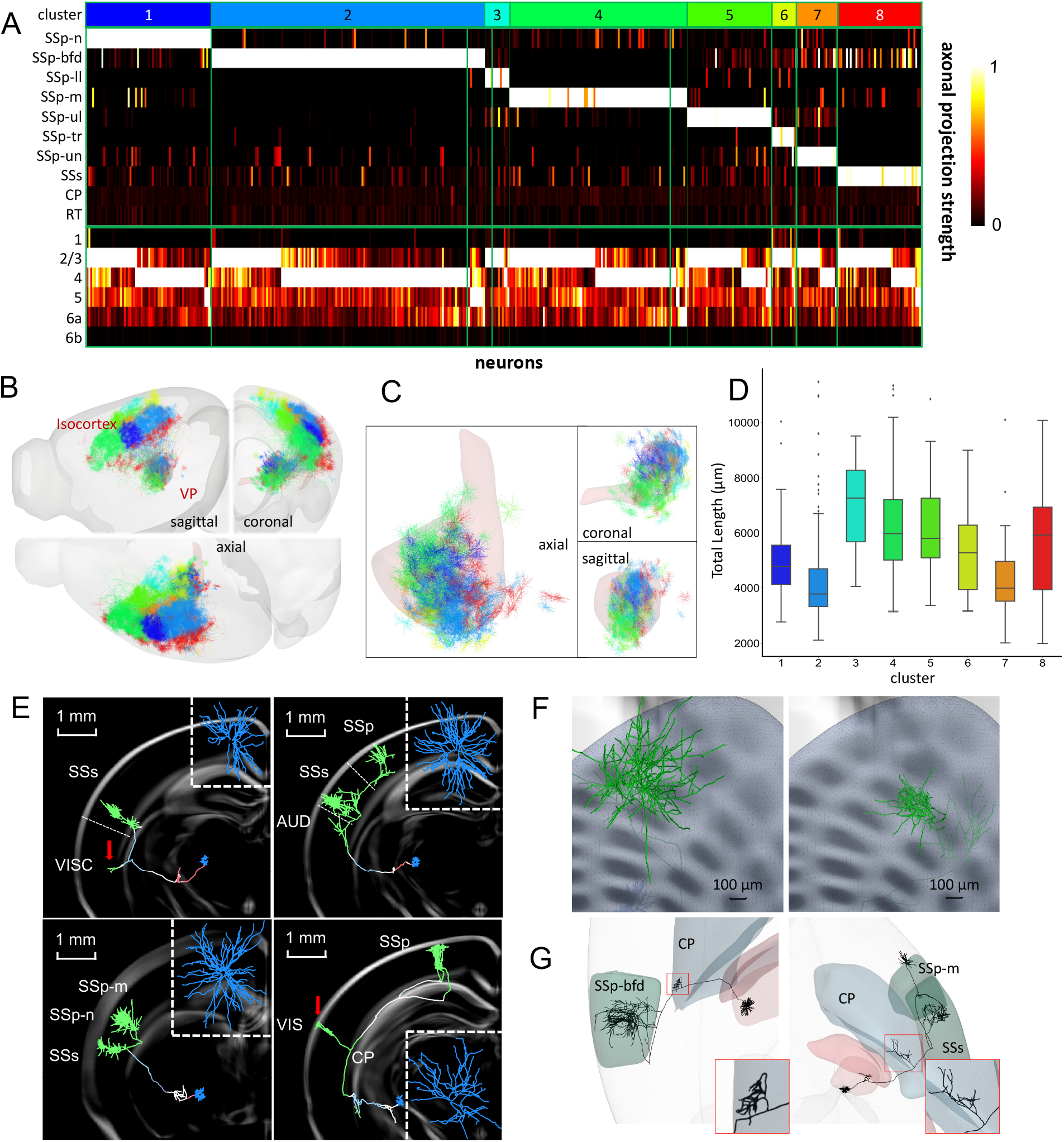
Alternative examination of connection types in ventral posterior (VP) nucleus. **A.** 8 different VP neuron subtypes clustered and color-coded by projecting target regions, particularly cortical layers (**Supplementary Table 8**). Columns: individual neurons. Rows: projection targets of neurons. Color bar: axonal length of a neuron projecting to a specific area. **B.** Axonal clusters of these VP subtypes mapped to CCF. **C.** Soma positions and connected dendrites of the 8 subtypes. **D.** Analysis of dendritic total length (μm) of these 8 clusters of neurons. **E.** Examples of VP neurons with zoomed-in coronal view of dendrites. Red arrows: projection targets outside of somatosensory areas; note the VIS target is in layer 1. **F.** Examples of cluster size located in the barrel field. Note the right cluster covers more than one barrel. **G.** Visualized CP projections of VPM neurons.

Furthermore, we found that a single VP neuron could target multiple sensory areas. For example, a VPM neuron can simultaneously projects to SSs and sub-areas of SSp, such as SSp-m, SSp-n, SSp-ll or SSp-ul (Figure 7A and **7E**). Indeed, some VPL neurons even project to layer 1 in addition to layer 4 (lower right panel in Figure 7E). Such neurons carry two separate axonal clusters: a larger one projecting to VPL neuron’s typical projection target, i.e., SSp-layer 4, and a smaller one targeting layer 1 of a different cortical area, such as SSs or even VIS (visual cortex).

Also of note, the surface-area of some VPM cells’ axonal cluster in SSp-bfd (largest: 384,942μm^2^) is twice larger than that of a barrel (Figure 7F). Traditionally, it is believed that each VPM cell only projects to one barrel (Pierret, et al, 2000). Our finding suggests potential signal regulation across multiple barrels; thus, the tactile sense signaling transmission could be a multithread process.

Additionally, 18.6% of VP neurons (*n* = 88) possess small branches with bouton terminations in subcortical striatum, suggesting VP-striatum projections (Figure 7G). Our finding indicates a new pathway in thalamic-subcortical circuit, supplementing the main pathway of VP nuclei to somatosensory cortex. Taken together, these single-cell VP reconstructions give clues to supplementary and complex signal transmission paths in multisensory integration circuits.

### Subtyping Target Connections of Thalamocortical Neurons in MOp Cortex

In addition to the outgoing “forward” connection patterns examined in preceding sections, we also investigated the diversity of incoming connections of a target brain region. Previous literature shows that the primary motor cortex (MOp) receives thalamocortical projection from the sensory-motor relay nuclei VAL and the modulatory or high order nuclei like VM or PO (Kuramoto, et al, 2011; Guo, et al, 2018; Guo, et al, 2000). Our analysis revealed additional connections from sensory relay nuclei VPM and VPL (Figure 7 and Figure 8).

With the whole-brain mapped full reconstructions we produced, it can be seen that individual neurons from PO, VM, VAL project to motor and somatosensory areas as a whole spectrum of connectivity subtypes (Figure 8A and **8B**). Indeed, projections of individual neurons display different layer preferences in thalamocortical areas. Such preference in MOp can be summarized as the following: PO neurons (n=14) focus on mainly layer 2/3 (4/14), layer 2/3 and layer 5 (4/14) and layer 5 (6/14). VAL neurons (n=34) have 5 main subtypes of connectivity projecting to (a) layer 1 (4/34), (b) layer 2/3 mainly (6/34), (c) layer 2/3 and layer 5 combined (6/34), (d) layer 5 mainly and layer 6 weakly (16/34), and (e) mainly layer 6a (2/34). VM neurons (n=13) have several subtypes projecting to layer 1 (8/13) combined with weak projection into other layers, layer 2/3 and layer 5 (4/13) and all layers (1/13). VP neurons (n=35) can be classified as subtypes including projections to layer 2/3 (30/35), layer 5 (3/35) and layer 6 (2/35), respectively (Figure 8B). Individual examples display axonal cluster phenotypes and projections (Figure 8C). Taken together, these new layer projections from individual thalamocortical neurons suggest fine regulations of the sensory-motor signal circuits.

**Fig 8.**
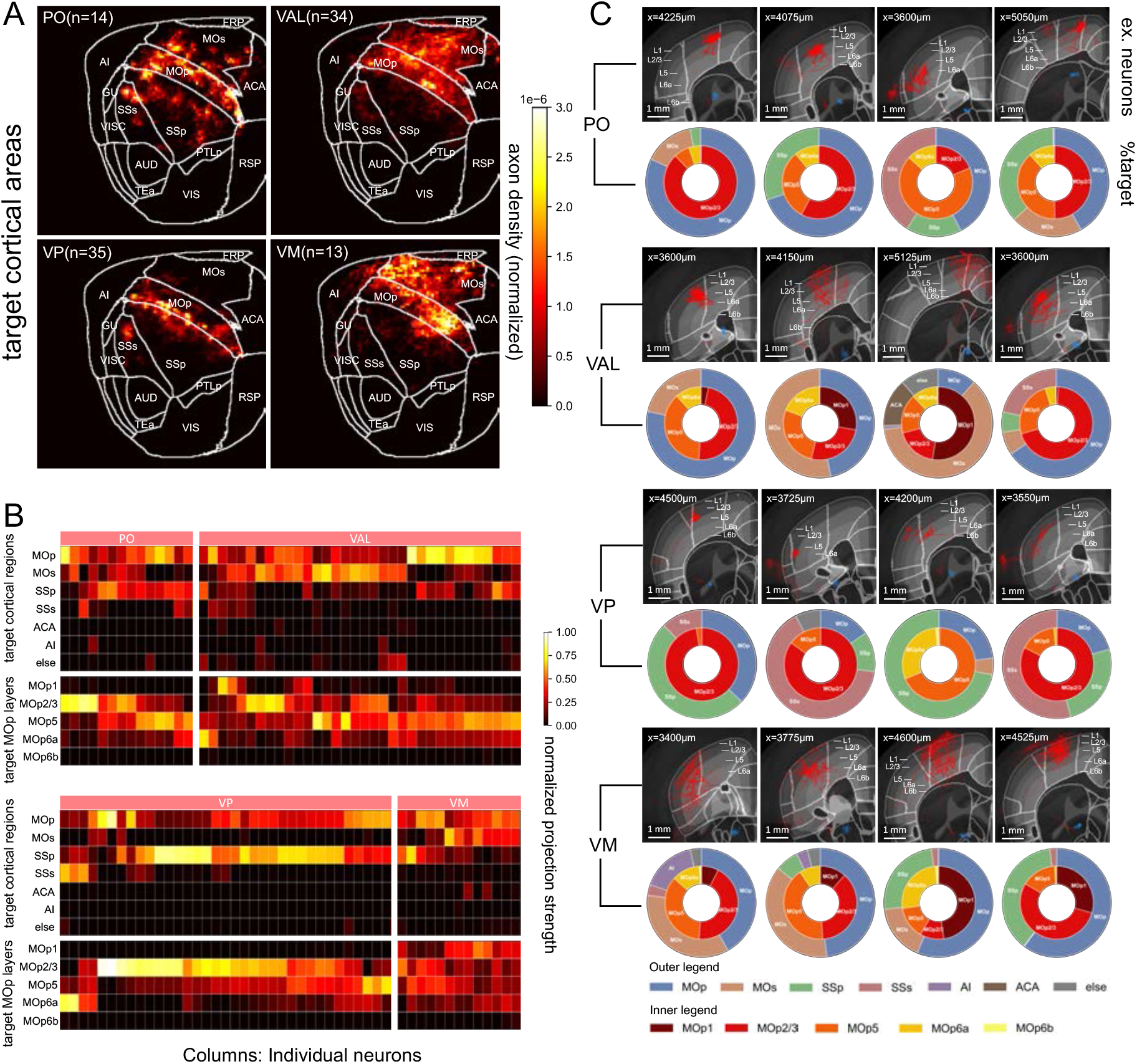
Conjugated MOp projections of individual thalamocortical neurons from PO, VM, VAL and VP nuclei. **A.** Co-projection of axonal arbors to MOp and nearby cortical areas originated from PO, VAL, VP, and VM. Color code: normalized arbor density. **B.** Projection matrices of individual neurons in **A**. Columns: individual neurons. Rows: projection targets, particularly with MOp layers. **C.** Example-neurons from each nucleus. Circular plots: distribution of target projection regions for each neuron.

## Discussion

This work studies the whole-brain scale connectivity of single neurons using one of the largest data archives produced to date, leveraging both new dendritic reconstructions that cover the entire brain and existing axonal and full reconstructions (Winnubst, et al., 2019; Peng, et al, 2021; Gao, et al, 2022). While multi-dataset aggregation enables powerful analysis, it also necessitates that these three existing datasets (Winnubst, et al., 2019; Peng, et al, 2021; Gao, et al, 2022) share cross-validated distributions of morphological properties of single neurons indicating quality reconstructions. The consistency of these data suggests a novel approach to reveal how individual neurons in different regions are wired into different networks with different circuit motifs at whole brain scale. There are two remarkable topics in such an integrative approach. First, one may be able to study the building blocks of a brain, i.e., organizational “types” or “subtypes” of individual neurons, in terms of connectivity. This work makes an initial attempt toward this end. Second, one may be able to construct and study the “microscale” connectome based on individual neurons, filling a gap between previous work at the population level mesoscale connectome and the nanoscale connectome that relies on using electron microscopy and/or other super-resolution microscopy methods more suitable for examining synaptic level connections of neurons in potentially smaller, local brain regions. We have also attempted the second approach in another ongoing study (unpublished work).

To understand the potential connectivity of neurons throughout a brain at single neuron resolution, it is essential to analyze the arborization of axons and dendrites in different anatomical areas. An overall axonal arborization distribution map (Figure 1) provides an understanding of the marginal distribution of neurons that innervate from different regions, and also highlights that arbors can be powerful entities to study neuronal connectivity. To complete this paradigm, we produced brain-wide dendritic arbor domains, which were used to generate the connectivity profiles for each individual neuron. In this way, the connectivity features can be precisely defined and utilized for analysis. This approach therefore constitutes a contribution essential for whole brain scale single-neuron analysis.

Our framework derives potential connectivity from full morphology plus atlas mapping into a standard space, so that the regional connection relationship of multiple neurons can be compared objectively with the appropriate context of their relative locations, distributions of their shapes, and spatial adjacency and/or overlap of their axon-dendritic arbors. Atlas mapping thus enables the expansion of previous approaches to neuron-type circuit analysis that were limited to local anatomical domains (Tecuatl, et al, 2021) to the entire mouse brain and long-range projection neurons. Our data show that connectivity features of neurons not only provide additional dimensions to distinguish neurons from different anatomical regions, but also allow effective neuron-typing when they are used alone. Within each of the anatomically established brain regions, we also see a strong correlation between the connectivity-based similarity and the spatial adjacency of neurons and their somas. Therefore, to approach the more challenging task of subtyping neurons, we can aggregate both connectivity and spatial information to observe distinct neuron-groups that are otherwise difficult to distinguish. Our application in analyzing MOs neurons demonstrates diversity of such neuron subtypes that cannot be readily inferred from existing data of neuron population-projections and molecular profiling. Further screening of the enriched regional connectivity of neurons in VP nuclei and MOp may provide additional evidence that connectivity subtypes do exist, and carry biological significance in signal relay and integration.

Within a general framework of cell typing, our study demonstrates that morphology cannot accomplish this alone. Indeed, based on this study and also previous work (Peng, et al, 2021), as well as converging results from invertebrate nervous systems (Mehta, et al, 2023), we hypothesize that t-types or e-types alone may also be insufficient, and it is an interesting open question how to synergize all these data in a common connectivity framework. We believe that there are two key steps to address this challenge. The first is the generation of connectivity associated t-type, e-type, and m-types data. A second step is building a thorough statistical model of all such data to mine the associations and distribution patterns, which could be homogeneous clusters or globally nonlinear manifold patterns (Liu and Qian, 2022).

This approach of leveraging connectivity-type analysis toward the determination and validation of neuronal cell types is powerful. It can be extended to brain-scale analysis of single neurons’ synaptic connectivity when data becomes available. An excellent example can be seen in the single-cell connectivity-types defined for a *Drosophila* brain (Scheffer, et al, 2020) that elaborate on the connection detail built upon morphological and lineage similarities. While such an approach provides the electron microscopy-based, ultrascale spatial resolution to precisely pinpoint synaptic connections, it is also subject to noise and imperfect process of data acquisition and computation, which would likely be exacerbated when applied to a much larger and complicated mammalian brain. The strength of the present approach is that we can readily study cell typing and subtyping using the arborization based regional connectivity, without precise pinpointing of synaptic level connections. This may be valuable when considering that individual synapses are subject to turnover via structural plasticity, while arbor geometry provides a relatively more stable circuit scaffolding (Stepanyants et al, 2002). Connection types and subtypes can also provide a useful blueprint of future synaptic level analysis. In summary, neuronal connectivity in mammalian brain provides a powerful discriminant in the classification of neuronal cell types, refining and adding novel class information to existing and widely studied modalities.

We caution that the subtlety in definitions of “morphology” and ‘connectivity” might cause slight confusion in a specific context. In literature, sometimes the analysis of morphology type might have used part of the connectivity information, such as the orientation-aligned neurons could be analyzed using lamination information (Gouwens, et a, 2019). The goal of this study, however, is to factorize the analysis in an understandable way. In this sense, our paradigm can contribute to a more organized and clear communication in this field.

## Acknowledgment

This project was mainly supported by a Southeast University initiative for neuroscience awarded to HP. HP was also supported by a Zhejiang Lab BioBit Program visiting grant (2022BCF07).

The Southeast University team was also supported by a MOST (China) Brain Research Project 2022ZD0205200/2022ZD0205204 awarded to LL. GAA was supported in part by NIH grants R01NS39600 and RF1MH128693. HZ was supported by BRAIN Initiative grant U19MH114830. We thank Sebastian Seung for comment and suggestion on the manuscript and literature, Yong Yao, Rafael Yuste, David Van Essen and a number of other experts for various inspirations including discussions, comments, suggestions, and community events.

## Author Contribution

HP conceptualized and designed this study, and managed the entire project. LL executed the project and supervised ZY and FX in generating all results along with the assistance of LMG and HC. HD, MH, and HZ all advised on the project and manuscript. GAA advised on selection and analysis of independent reconstructions for validation, participated in critical discussion on the early phases of analysis design, and assisted in editing and revising the manuscript focusing specifically on relation with extant scientific literature. HP wrote the manuscript with input from all coauthors.

## Competing interests

All other authors declare they have no competing interests.

## Data and materials availability

The Supplementary Tables can be found along with the submission files of this manuscript. The key data tables of the analysis are currently available at https://braintell.org/projects/mousectype/Materials_0516.zip. The local reconstructions of dendrites are available at https://braintell.org/projects/mousectype/SEU_10860_dendrite.zip.

## Methods

### Full and axon reconstructions

We performed a detailed analysis of 1741 fully reconstructed single neuron morphologies (Peng, et al, 2021) called BICCN AIBS/SEU-ALLEN, and 1200 full single neuron reconstructions from the Janelia MouseLight project (Winnubst, et al, 2019). We also analyzed the axonal morphology of 6357 neurons generated by ION (Gao, et al, 2022). All neurons were registered to the Allen Mouse Brain Common Coordinate Framework v3 (CCFv3). These data are also documented in **Supplementary Table 1.** The naming convention of brain regions follows the CCFv3 and also consistent with the previous studies. Abbreviations are also recorded in **Supplementary Table 2.**

### Generation of dendritic tracing

We generated 10860 dendrite reconstructions from fMOST imaging with the following protocol. First, we collected image samples following the same protocol in our previous study on generating the full reconstructions (Peng, et al, 2021). Next, we ran the APP2 algorithm (Xiao and Peng, 2013) for tracing local arbors by taking manually defined and validated somas as the central starting points in local image volumes (1024×1024×512 voxels), for the goal that the joint area of these local volumes covers main dendrite arbors. We ran APP2 with a number of background thresholds (10,15, 20, 25, 30, 35) resulting in 6 tracing candidates. Then, we leveraged the set of manually annotated and validated dendritic arbors (from MouseLight and BICCN AIBS/SEU-ALLEN) to filter the automatic tracing results. The [min, max] interval of the following five features of the dendritic arbors were considered realistic, including ‘Tips’ [7, 143], ‘Length’ [700, 13615], ‘Max Path Distance’ [108, 1382], ‘Average Bifurcation Angle Remote’ [35, 129], and ‘Max Branch Order’ [3, 32]. An automatic tracing would be discarded if less than four of its features fell out of these limits. In case more than one tracing qualified for a soma location, we kept the tracing with greatest length. We spatially registered all tracings to CCFv3 using mBrainAligner (Qu, et al, 2022). In total, we collected images from 53 mouse brains, we identified 31,625 neurons, and we generated 17,228 qualified tracings. We visually inspected all tracings and discarded those with obvious errors (e.g., 1 trace covers multiple touching neurons, mis-alignment during registration), finally obtaining 10,860 proofread dendritic tracings.

### Independent reconstructions for validation

To cross-validate the neuron morphologies used in this work, we also considered independent morphologies produced and documented in public resources. Particularly, we searched adult mouse neuron reconstructions in certain brain regions via keywords using the searching tool of NeuroMorpho.Org (http://neuromorpho.org/KeywordSearch.jsp). When possible, we only kept the neuron reconstructions tagged as complete or, in case those were not available, moderately complete. We searched neurons in 4 different brain regions (HPF, SS, MO and ACA; see **Supplementary Table 2** for a complete list of abbreviations) using keywords “{region}&dendrite&mouse&adult” where {region} was one of the 4 acronyms. Details of data sources are listed in **Supplementary Table 3** (Yamashita, et al, 2018; Gong, et al, 2016; Iascone, et al, 2020; Cohen, et al, 2013; Smit-Rigter, et al, 2012; Suter, et al, 2015; Jiang, et al, 2020; Lin, et al, 2018; Morelli, et al, 2014; Murase, et al, 2016; Karlsson, et al, 2016).

### Gaussian Mixture Model classification of neuron nodes

We resampled fully traced SWC files (*n* = 9298) to have nodes every 10 μm and saved their coordinates in the space of the Common Coordinate Framework v3 (CCFv3) (Wang, et al, 2020) at 25μm isotropic voxel resolution. For each of the 19 CCFv3 brain regions with most neurons in the analyzed datasets (MOs, AId, ACAd, ACAv, ORBvl, ORBl, ORBm, VPM, CP, AIv, FRP, ILA, MOp, SSp, VPL, SUB, LGd, SSs; see **Supplementary Table 2** for reference), we pooled all SWC coordinates in a single data frame containing their x, y, and z locations. We clustered the pooled data using the Mclust function with default parameters (mclust R package version 5.4.7 (Scrucca, et al, 2016)). We selected the Gaussian Mixture Model (GMM; among all combinations of spherical, diagonal and ellipsoidal with equal or varying volume, shape and orientation) as it provides optimal clustering as measured by the Bayesian Information Criterion (BIC) (Schwarz, 1978). BIC is a measure for the comparative evaluation among a finite set of statistical models, based on maximizing the likelihood function while penalizing for the number of parameters in the models. We saved the resulting classification with the node IDs of each neuron.

### Definition of arbor domains using α-shape

We defined 3-D dendritic domains by using the pooled, clustered SWC coordinate dataset. We found the minimal volume enclosing all nodes belonging to each single cluster by obtaining the 3D α-shape of the point set (alphashape3d R package version 1.3.1) (Edelsbrunner, et al, 1994). The 3D α-shape is a generalized definition derived from the Delaunay triangulation (Delaunay, et al, 1934) with a parameter α to control for the level of detail (the convex hull is obtained when α~ꝏ). To obtain detailed volumes enclosing all neuron nodes in each cluster, we used α=0.4. We call the obtained 3D shapes “arbor domains”. When the majority of the nodes within an arbor domain belonged to neurons with their soma in the domain itself, we categorized those as dendritic arbor domains. Otherwise, we considered the obtained domains to be axonal. We saved all arbor domains as surface objects. We plotted 2D slices of the arbor domains using the R base plot function (version 4.1.0).

The definition of dendritic domains based on full tracings was obtained both for raw data distributed in both brain hemispheres and for flipped neurons, ensuring that all of them had somas in the same hemisphere. To further analyze connectivity, in that case, dendritic domains were flipped to also recapitulate homologous contra-lateral regions.

In addition to the arbor domains obtained from fully traced neurons, we also generated single dendritic domains from using all node coordinates for dendritic tracings with somas inside each of the 19 brain regions with most neurons. 3D α-shapes were defined using the same method. However, in this case we did not perform GMM-clustering and all coordinates were pooled in a single set for each brain hemisphere.

### Single neuron connectivity to dendritic arbor domains

To define outgoing connections from single fully traced neurons to dendritic arbor domains, we measured the spatial overlap between single neurons and arbor domains. To do so, we obtained all voxels enclosed by each domain 3D α-shape (we tested whether they are inside the surface of the domain using the inashape3d function in the alphashape3d R package version 1.3.1) and saved them as a 3D mask in the CCFv3 space. To convert surface polygon file format .ply files to 3D masks we used binvox version 1.35 (Nooruddin, et al, 2003). We then obtained a 3D volume where each voxel contains an array of indices identifying each 3D α-shape volume visiting such voxel. We obtained α-shapes for each individual fully traced neuron (α=0.4) and saved the enclosed volume as a 3D mask. Finally, we measured the overlap volume between each single neuron mask and the volume containing all 3D arbor domain indices. We saved the overlapping volume between each neuron and each dendritic domain as a connectivity matrix.

### Support Vector Machine clustering of morphology and connectivity

To assess the relevance of arbor domain connectivity for defining cell types and subtypes, we used a Support Vector Machine (SVM; hyperoverlap R package version 1.1.1; linear kernel, cost=1000 and stoppage.threshold=0.2) to classify neurons with somas located in MOp, SUB and VPL regions (Brown, et al, 2020; Cortes, et al, 1995). For each pair of brain regions, we used SVM to cluster the data in two groups. To assess the separation of the neurons in the space defined by the two morphological variables “total length” and “maximum branch order”, we measured the pairwise overlap of points from each of the three brain regions. To account for arbor domain connectivity, we obtained a PCA from the connectivity matrix of the analyzed neurons. We performed pairwise SVM classification analogously by adding the first three principal components of the connectivity matrix in the dataset. We plotted these results using the ggplot2 R package (version 3.4.0).

### m/c-score metric

The m/c-score can be used to quantify the dissimilarity of morphological (see **Supplementary Table 5** for axonal features and **Supplementary Table 6** for dendritic features) and connectivity (arrays of spatial overlap between each single neuron and all dendritic arbor domains) features between two clusters, taking into account both their intra-class similarity and inter-class separation. A higher score indicates the greater difference between two clusters, while a lower score indicates more similarity. The m/c-score is calculated as the following formula:

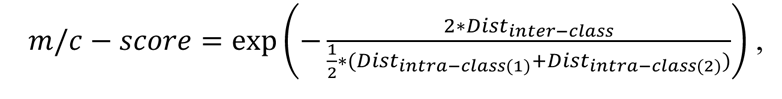

where, *Dist*_inter-class_represents inter-class distance between the centers of two clusters, which is calculated using Manhattan distance metric (Han, et al, 2022). *Dist*_*intra-class(x)*_represents intra-class distance of cluster *x*, which is defined as the average of the Manhattan distances between each sample and all other samples within the same cluster.

With regard to m-score matrix clustering, we applied hierarchical clustering by clustermap function (method=“ward”, metric=“euclidean”) in Seaborn Python package (version 0.11.2). We used umap-learn Python package (version 0.5.1) to implement UMAP decomposition with default parameters and plotted results as scatterplots with Matplotlib (version 3.3.4).

### Anatomy-based distance metric

The distance metric follows the Mahalanobis definition (McLachlan, et al, 1999). Let *S*_*j*_ = [*x*_*j*_, *Y*_*j*_, *Z*_*j*_] be the position of soma *i* in 3-D space. Due to the computational convenience, the soma location should be mirrored to the ipsilateral hemisphere. For two somas *S*_1_, *S*_2_, anatomy-based distance was defined using the following equation:

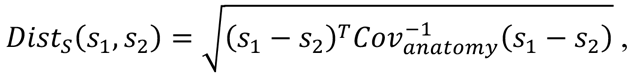

where, *Cov*_*anatomy*_ represents the covariance of 3D positions of voxels of relevant ipsilateral anatomical region in the 25μm CCFv3 reference space volume.

### Distance-weighted connectivity-based clustering

Distance-weighted connectivity-based clustering was used to cluster s-type cells based on both their connectivity feature similarity and physical distance of somas. Two matrices were generated to represent these components: a connectivity similarity matrix (c-similarity matrix denoted by *M*_*C*_) calculated using cosine similarity, and a distance matrix (d-map denoted by *M*_*D*_) calculated based on the anatomy-based distance between the somas of the cells. Both matrices were linearly normalized to values between 0 and 1. To emphasize spatial adjacency, a distance affinity matrix (*M*_*DA*_) was constructed using Gaussian kernel; *M_DA_* = exp(−*M_D_* · *M_D_*). This ensured that larger values in the affinity matrix indicated greater spatial adjacency between cells. Hierarchical clustering was subsequently applied on the matrix resulting from multiplying *M_C_*and *M_DA_* to produce diversity clustering results. The optimal number of clusters was determined by the Calinski-Harabasz (Caliński, et al, 1974) score (metrics.calinski_harabaz_score function from scikit-learn Python package version 0.24.2) automatically.

### Correlation between single cell and population morphology, projections, and transcriptomics

We used a transcriptomic dataset of 34,331 neurons in MOs and FRP brain regions, which is collected from a newly released dataset (Yao, et al., 2021). The analysis is performed by SCANPY (a python package, version: 1.9.3). To ensure the data quality, we filtered out 9432 genes that are detected in less than 3 cells, and filtered out 624 cells that expressed over 6,000 genes. We normalized the data (using functions: pp.normalize_total, pp.log1p, pp. regress_out, under default parameters), and reduced its dimension (using tl.pca, and tl.umap, under default parameters) for visualization. We further extracted L5 related cells (9,879 cells) using this genetic modality, following the same procedures.

### Electrophysiological data analysis

For electrophysiological modality, we selected 919 cells in VISp layers from a Path-seq dataset (Gouwens, et al., 2020). The selected dataset has 5 transcriptomic labels (Pvalb Reln Itm2a, Sst Hpse Cbln4, Sst Calb2 Pdlim5, Lamp5 Lsp1, Pvalb Sema3e Kank4), and 5 strcuture labels (VISp1, VISp2/3, VISp4, VISp5, VISp6a). UMAP layout of the dataset shows three distinct populations. For 919 cells in electrophysiological profile, we used IPFX (a python package, version: 1.0.7) for the feature extraction, generating 13 electrophysiological features (**Supplementary Table 7**) for each cell. We concatenated these features as one vector profiling each cell in subsequent analyses.

---

**Supplementary Figure 1.**
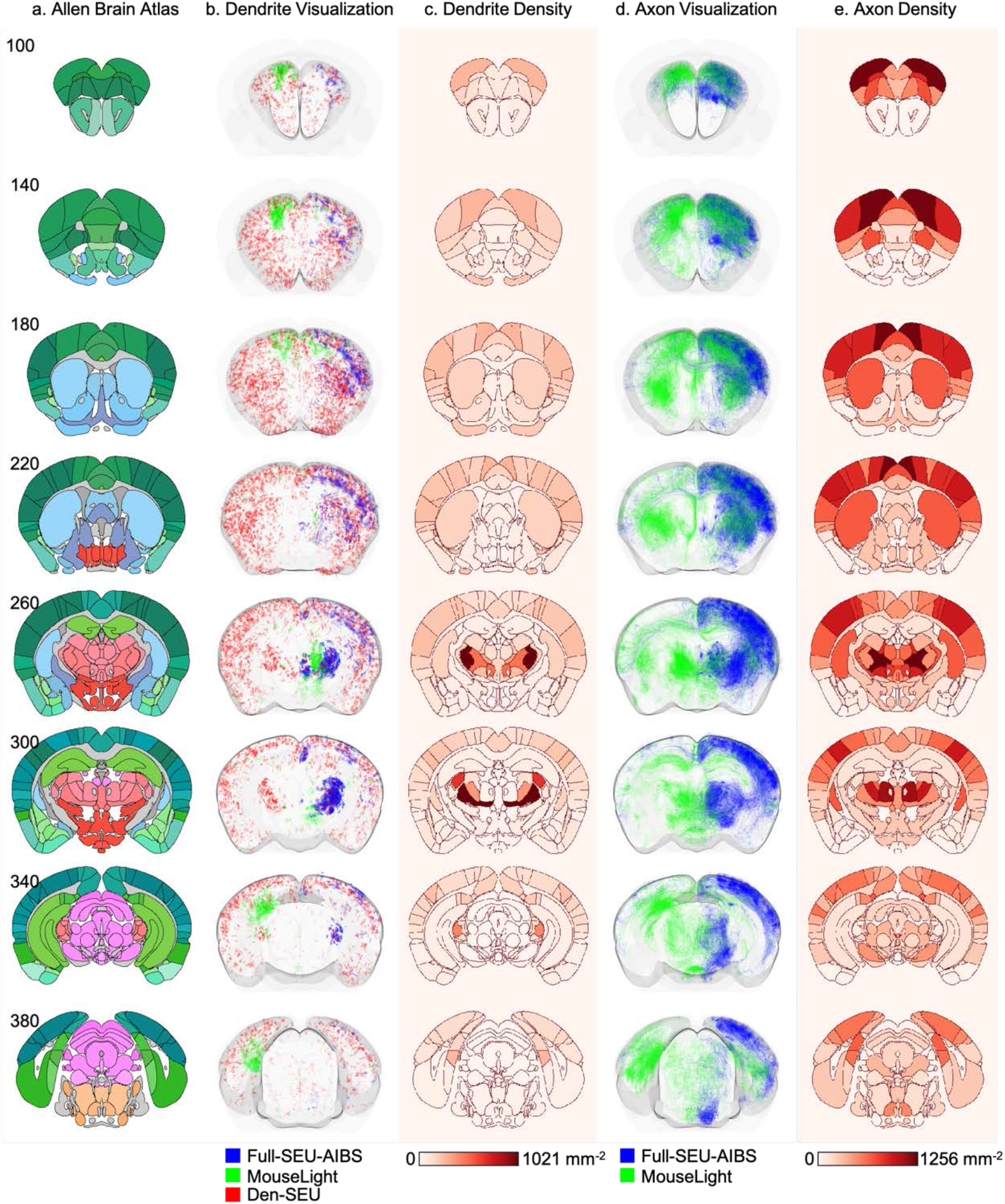
Exemplar summary of the spatial anatomical distribution of several neuron reconstruction datasets. Eight axial slices are selected for visualization. (a) CCF atlas showing brain regions of selected slices. Brain regions are colored following CCF’s color code. CCF slice ID is shown on the top-left of each image. (b) Visualization of dendrite reconstructions. (c) Dendrite density within each brain region. (d) Visualization of axonal arbor reconstructions. (e) Axonal arbor density within each brain region. (b) and (d), In each image, dendrite/axonal arbors within 500 *μm* (20 slices) of the target slice are shown. Different color is assigned to different dataset. (c) and (e), Density is computed by dividing total dendrite/axonal arbor length (mm) inside a brain region by the volumetric size (mm^3^) of the brain region. The unit of arbor density is mm^-2^. The color map is shown on the bottom.

**Supplemental Figure 2.**
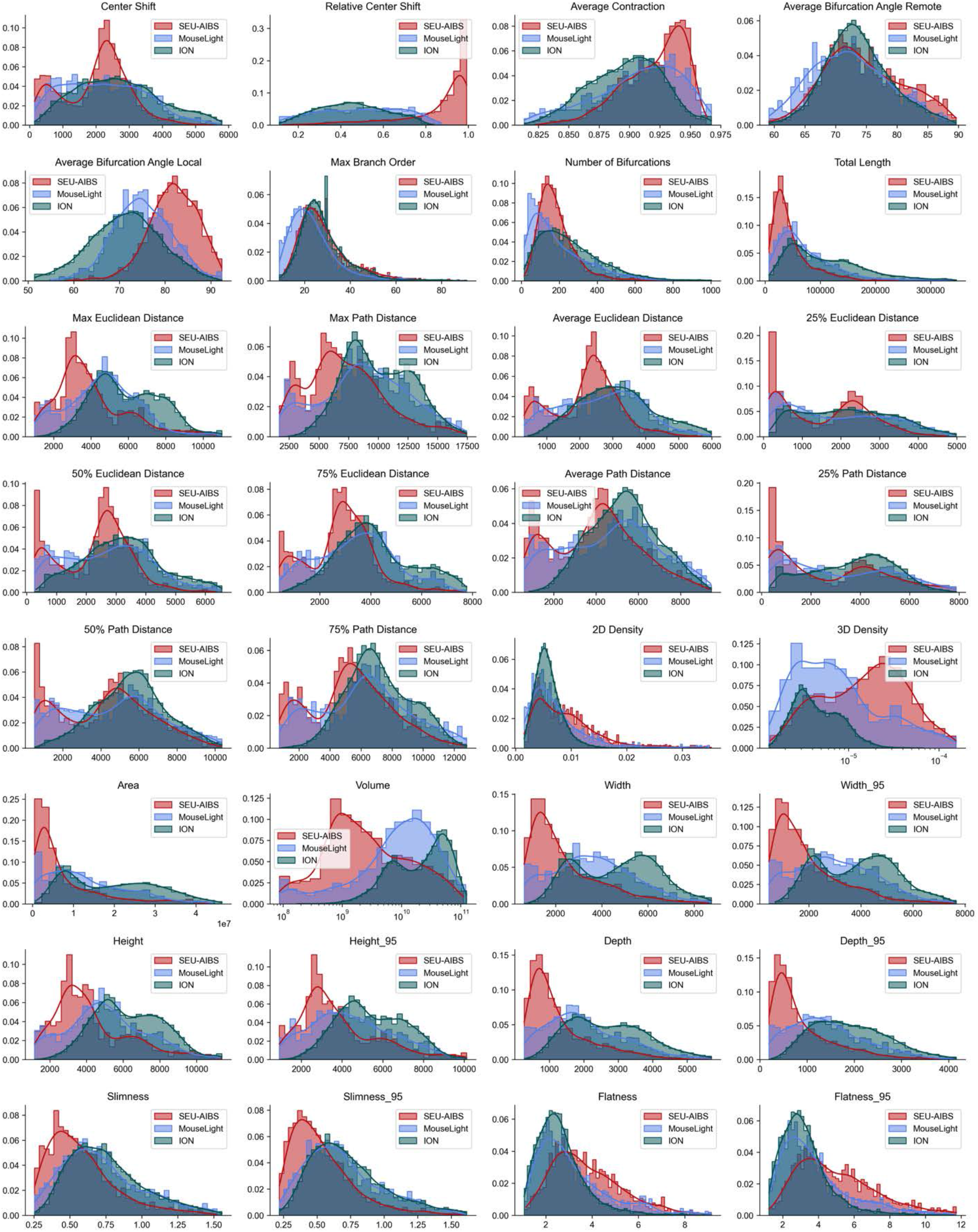
Comparative analysis of morphological features of axons in three datasets, i.e. BICCN AIBS/SEU-ALLEN (SEU-AIBS), Janleia MouseLight (MouseLight), and ION.

**Supplemental Figure 3.**
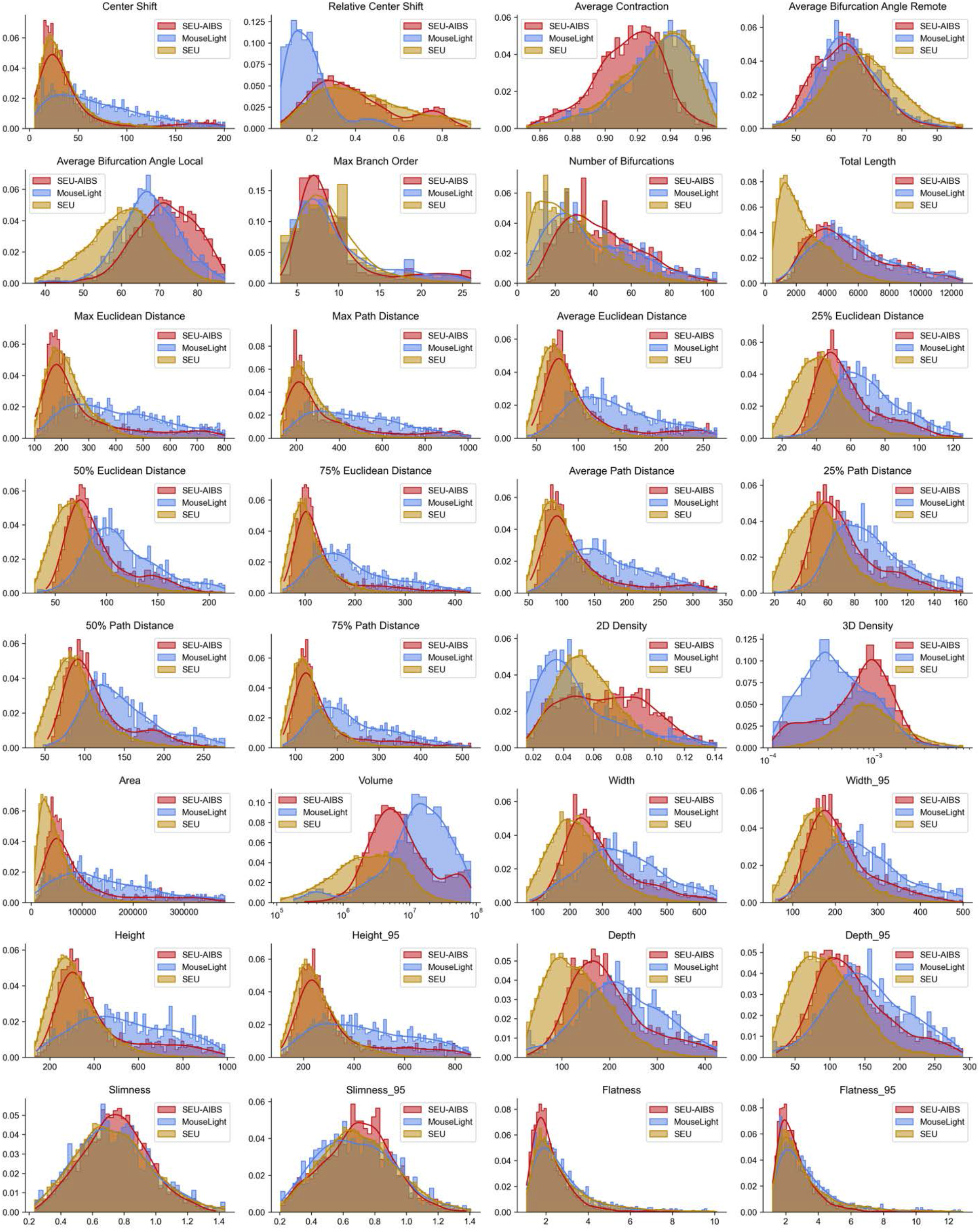
Comparative analysis of morphological features of dendrites in three datasets, i.e. BICCN AIBS/SEU-ALLEN (SEU-AIBS), Janleia MouseLight (MouseLight), and DEN-SEU (SEU).

**Supplemental Figure 4.**
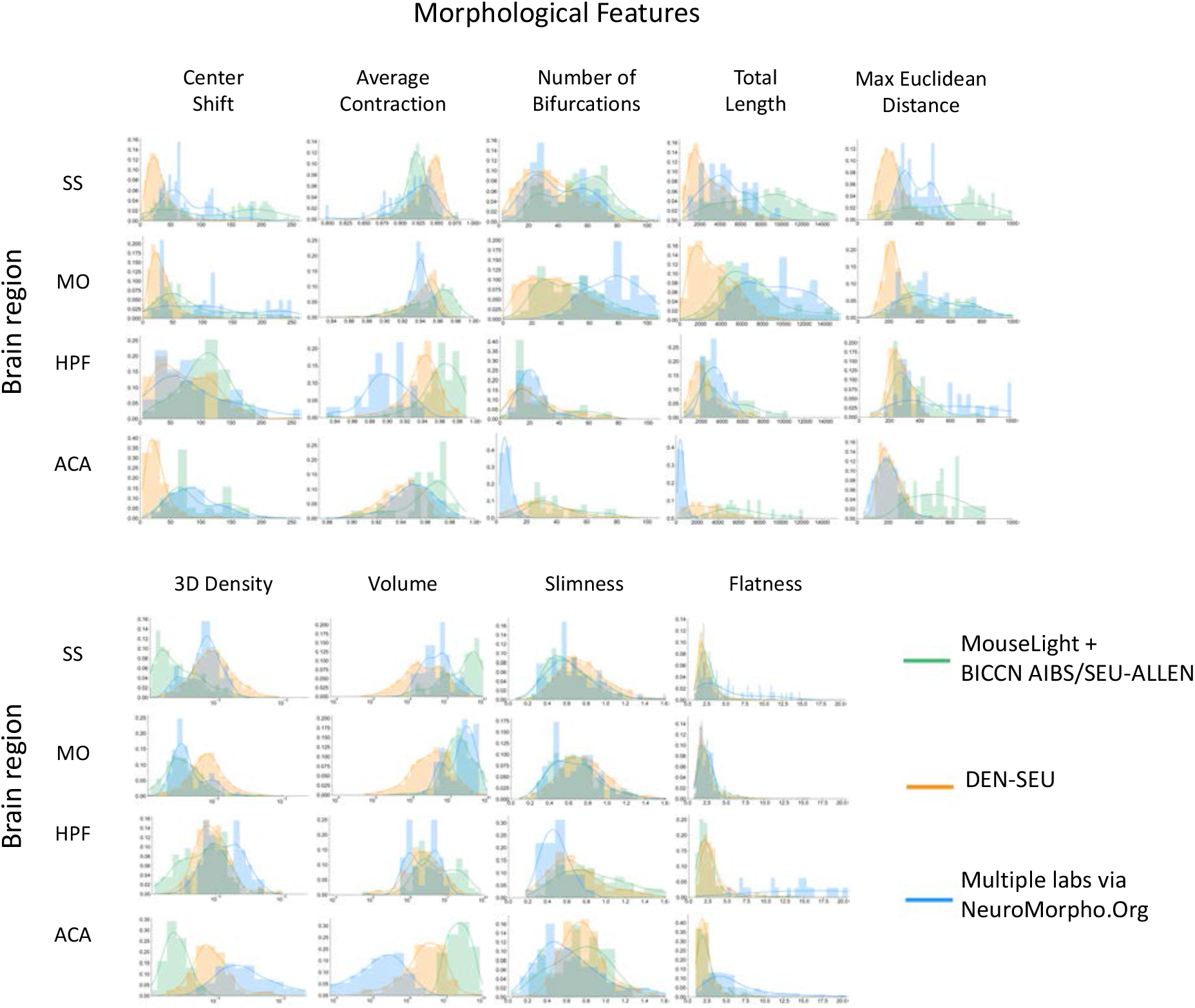
Comparative analysis of dendrite morphology features of neurons in selected brain regions, for multiple datasets including full single neuron reconstructions (BICCN AIBS/SEU-ALLEN and MouseLight), dendritic reconstructions (DEN-SEU), and publicly available reconstructions from multiple independent labs (as archived at NeuroMorpho.Org, see **Methods**). Each row corresponds to a brain region, while each column corresponds to a feature. Four brain regions, i.e. somatosensory area (SS), somatomotor area (MO), hippocampal area (HPF), and anterior cingulate area (ACA), with available neuron feature data were selected. Nine informative features are shown as examples. Refer to **Supplementary** Figure 5 for a complete comparison of all 32 features.

**Supplemental Figure 5.**
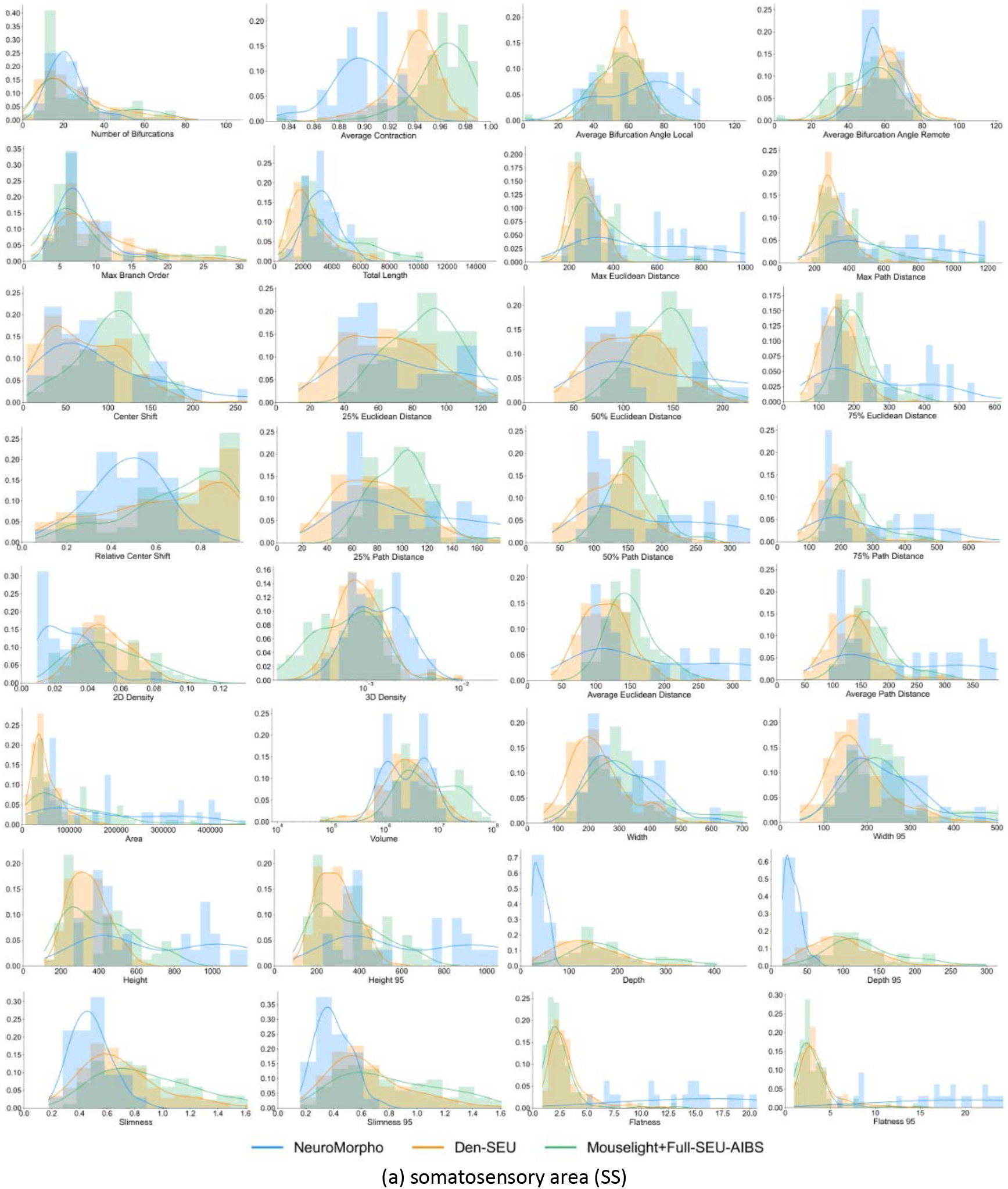
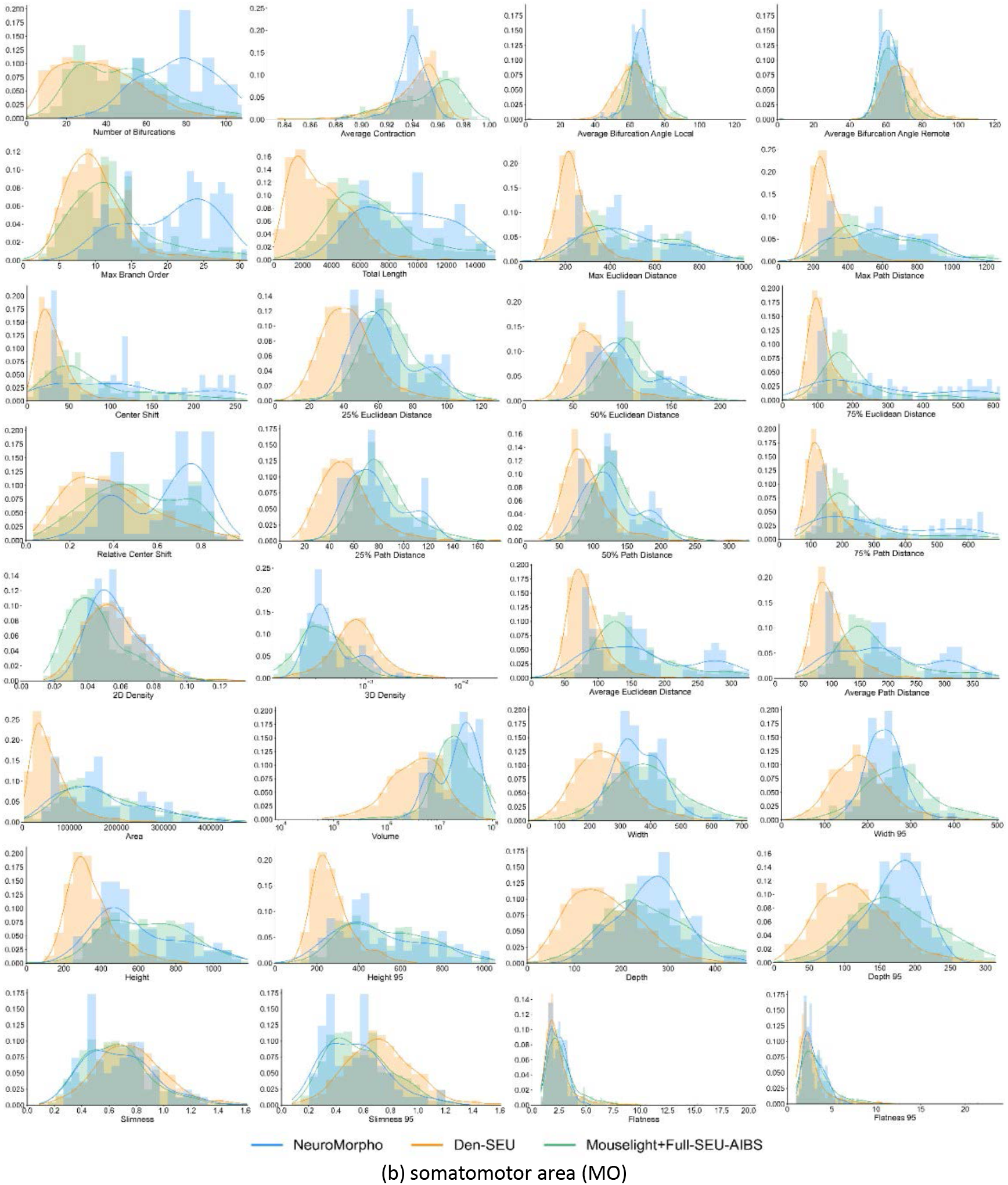
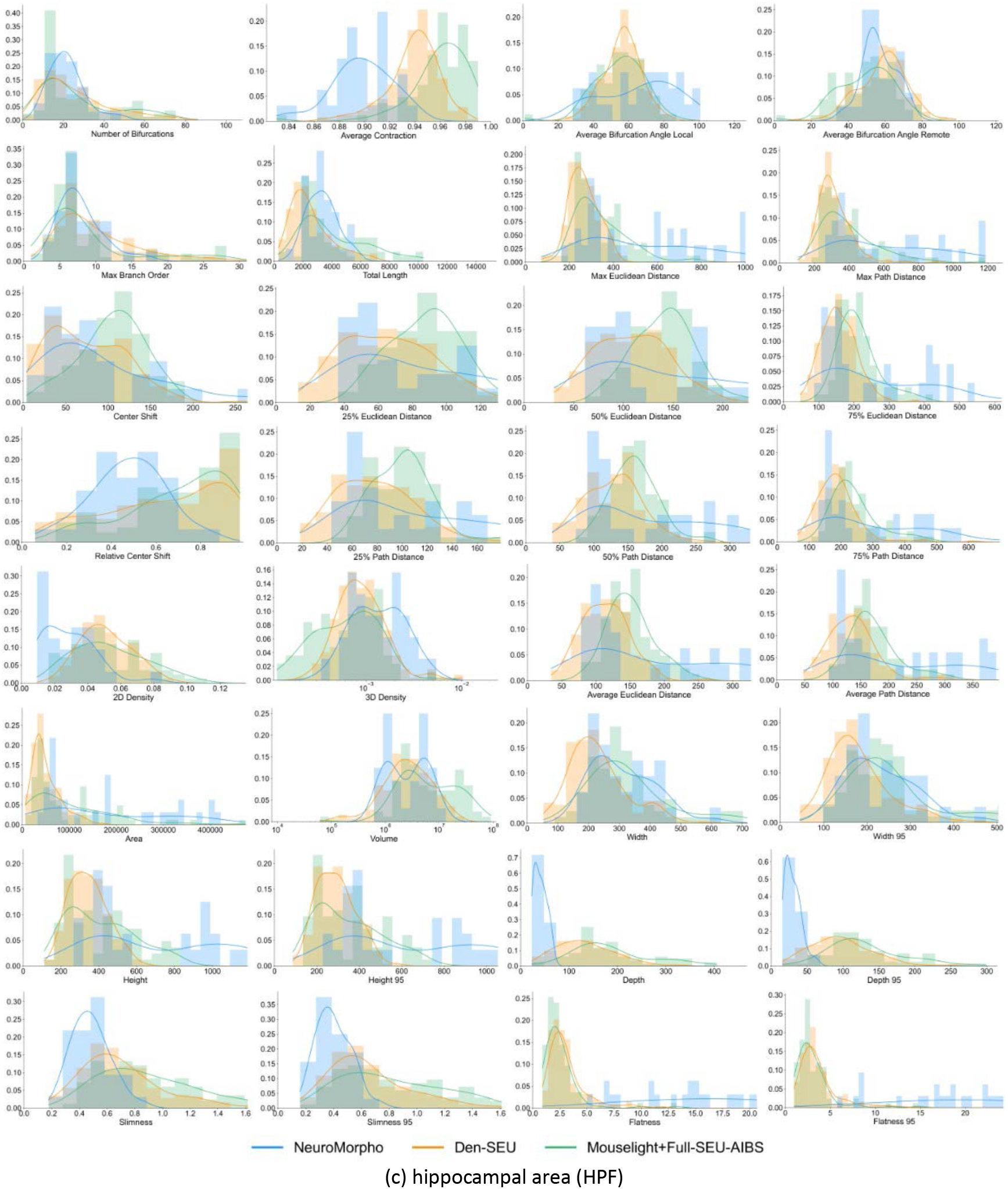
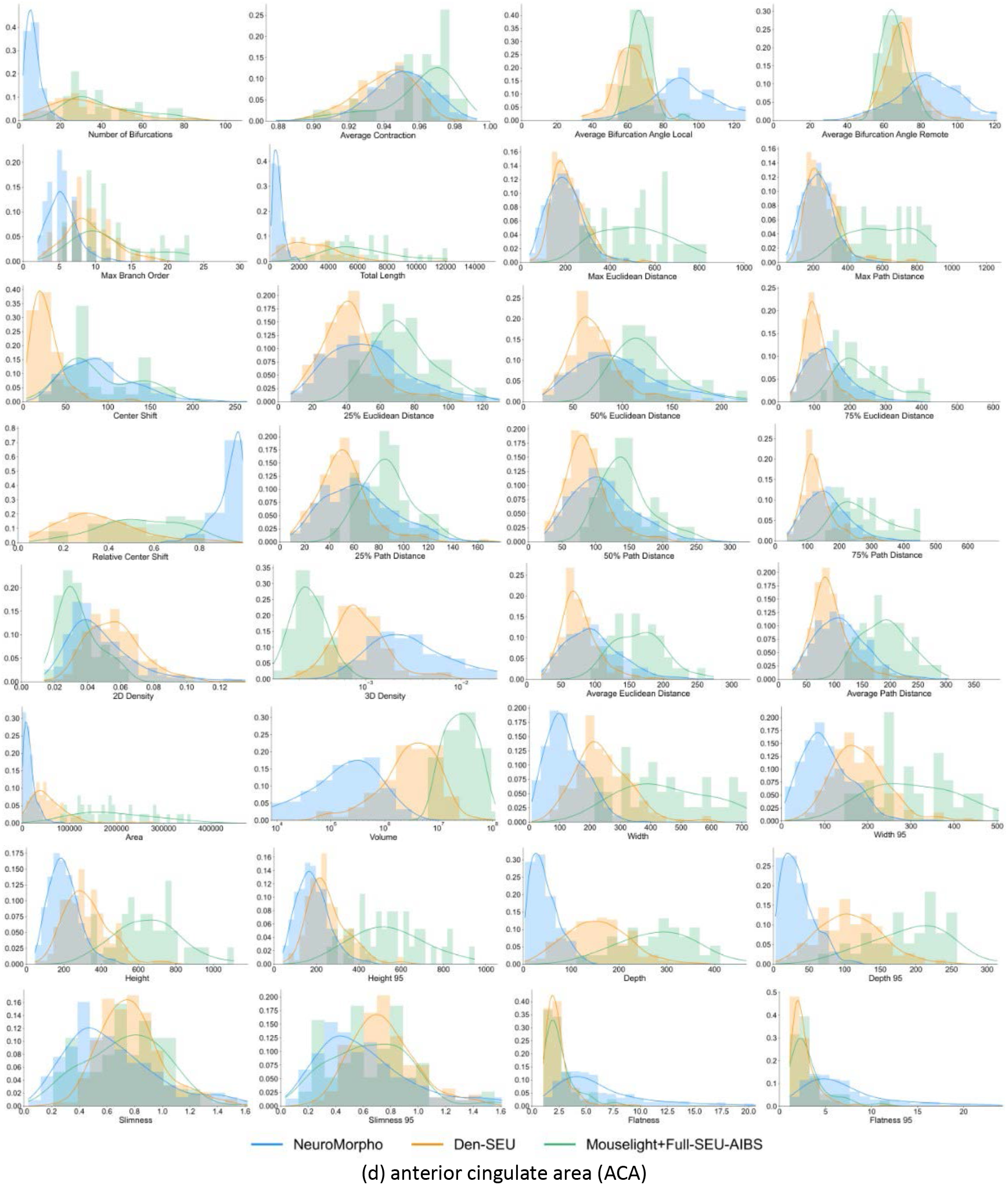
Comparative analysis of dendrite morphology features of neurons in selected brain regions, for multiple datasets including full single neuron reconstructions (BICCN AIBS/SEU-ALLEN and MouseLight, labeled as “MouseLight+Full-SEU-AIBS”), dendritic reconstructions (DEN-SEU/“Den-SEU”), and publicly available reconstructions from multiple independent labs (“NeuroMorpho”, as archived at NeuroMorpho.Org, see **Methods**). Four brain regions, i.e. (a) somatosensory area (SS), (b) somatomotor area (MO), (c) hippocampal area (HPF), and (d) anterior cingulate area (ACA), are shown with the comparison, in which all 32 morphology features are visualized with the respective names under each subplot.

**Supplemental Figure 6.**
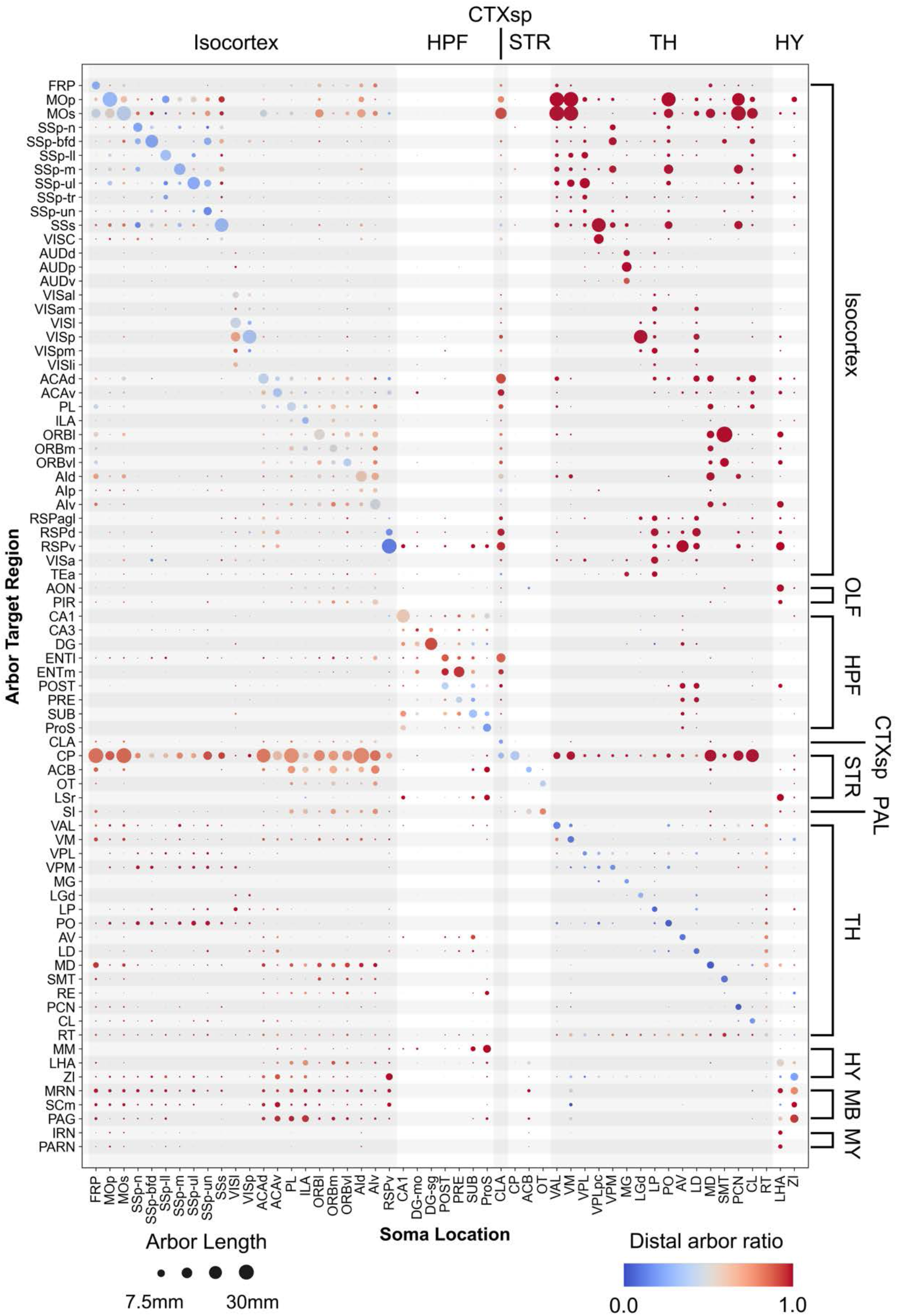
Whole brain arborization map of all neurons with axons in this study. See **Figure 1D** for labels of brain regions.

**Supplemental Figure 7.**
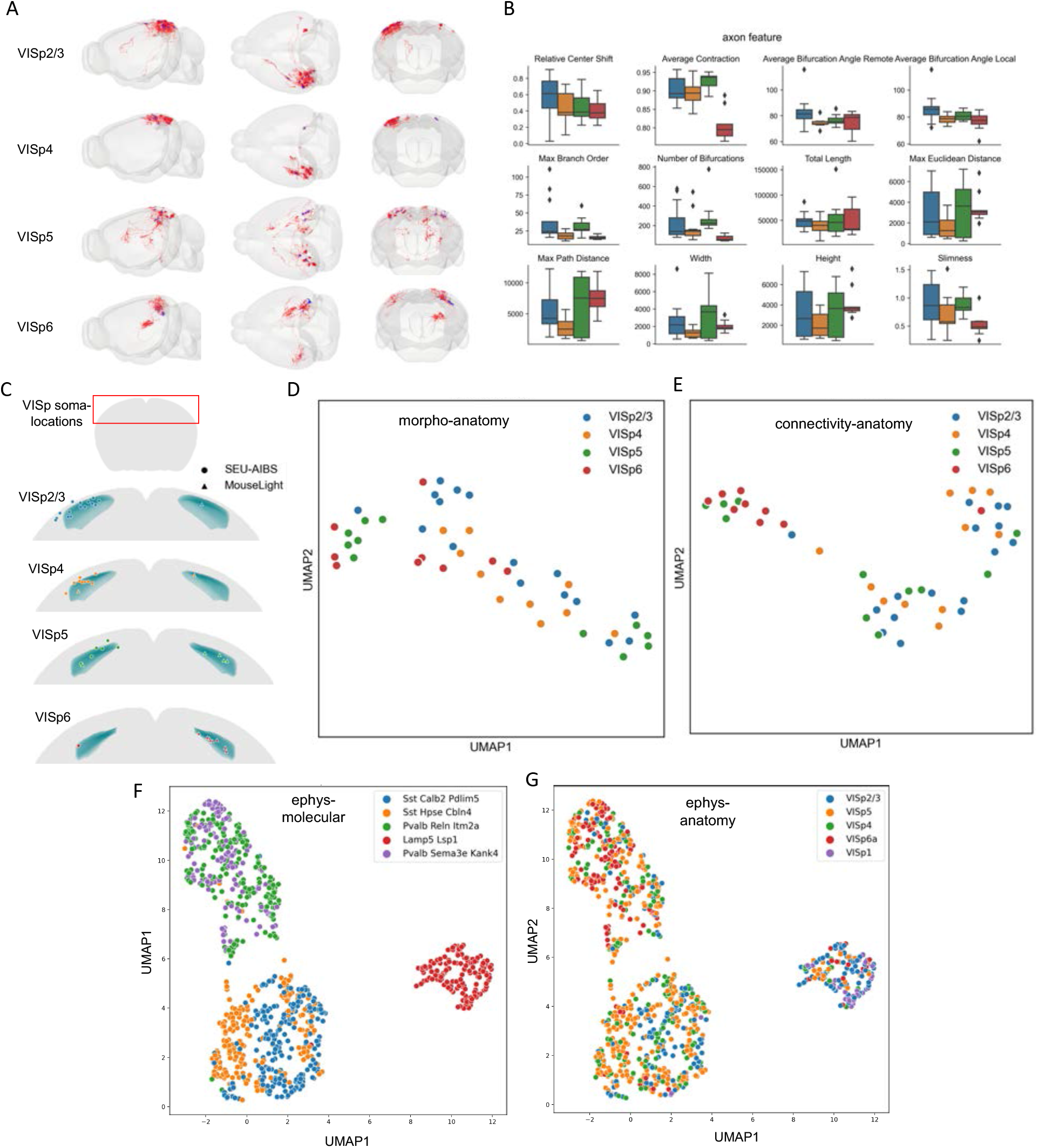
Comparative analysis of projection patterns of VISp neurons originated in various cortical layers (2/3, 4, 5, and 6), the respective morphological features and soma locations, and public-domain electrophysiological recording and transcriptomic profiles of single neurons. **A**. Projection and regional connectivity patterns of VISp neurons, grouped by soma-locations in four cortical layers. **B**. Comparison of axon features of VISp neurons in four layers. **C**. Locations of VISp neurons used in this study. **D**. Joint distribution of morphological features and soma locations in the respective UMAP space. **E**. Joint distribution of connectivity features and soma locations in the respective UMAP space. **F**. Joint distribution of electrophysiological features and molecular types of VISp neurons in the respective UMAP space. **G**. Joint distribution of electrophysiological features and soma locations in the respective UMAP space.

## Notes

### Competing Interest Statement

The authors have declared no competing interest.

## References

1. Abbott, L. F., Bock, D. D., Callaway, E. M., Denk, W., Dulac, C., Fairhall, A. L., … & Van Essen, D. C. (2020). The mind of a mouse. Cell, 182(6), 1372–1376.

2. Akram, M. A., Nanda, S., Maraver, P., Armañanzas, R., & Ascoli, G. A. (2018). An open repository for single-cell reconstructions of the brain forest. Scientific data, 5(1), 1–12.

3. Axer, M., & Amunts, K. (2022). Scale matters: The nested human connectome. Science, 378(6619), 500–504.

4. Bijari, K., Akram, M. A., & Ascoli, G. A. (2020). An open-source framework for neuroscience metadata management applied to digital reconstructions of neuronal morphology. Brain Informatics, 7(1), 1–12.

5. Binley, K. E., Ng, W. S., Tribble, J. R., Song, B., & Morgan, J. E. (2014). Sholl analysis: a quantitative comparison of semi-automated methods. Journal of Neuroscience Methods, 225, 65–70.

6. Bureau, I., von Saint Paul, F., & Svoboda, K. (2006). Interdigitated paralemniscal and lemniscal pathways in the mouse barrel cortex. PLoS Biology, 4(12), e382.

7. Clascá, F., Rubio-Garrido, P., & Jabaudon, D. (2012). Unveiling the diversity of thalamocortical neuron subtypes. European Journal of Neuroscience, 35(10), 1524–1532.

8. Cortes, C., & Vapnik, V. (1995). Support-vector networks. Machine Learning, 20, 273–297.

9. Dong, H. W. (2008). The Allen reference atlas: A digital color brain atlas of the C57Bl/6J male mouse. John Wiley & Sons Inc.

10. Dorkenwald, S., Turner, N. L., Macrina, T., Lee, K., Lu, R., Wu, J., … & Seung, H.S. (2022). Binary and analog variation of synapses between cortical pyramidal neurons. Elife, 11, e76120.

11. Gao, L., Liu, S., Gou, L., Hu, Y., Liu, Y., Deng, L., … & Yan, J. (2022). Single-neuron projectome of mouse prefrontal cortex. Nature Neuroscience, 25(4), 515–529.

12. Gouwens, N. W., Sorensen, S. A., Baftizadeh, F., Budzillo, A., Lee, B. R., Jarsky, T., … & Zeng, H. (2020). Integrated morphoelectric and transcriptomic classification of cortical GABAergic cells. Cell, 183(4), 935–953.

13. Guido, W. (2018). Development, form, and function of the mouse visual thalamus. Journal of neurophysiology, 120(1), 211–225.

14. Guo, K., Yamawaki, N., Barrett, J. M., Tapies, M., & Shepherd, G. M. (2020). Cortico-thalamo-cortical circuits of mouse forelimb S1 are organized primarily as recurrent loops. Journal of Neuroscience, 40(14), 2849–2858.

15. Guo, K., Yamawaki, N., Svoboda, K., & Shepherd, G. M. (2018). Anterolateral motor cortex connects with a medial subdivision of ventromedial thalamus through cell type-specific circuits, forming an excitatory thalamo-cortico-thalamic loop via layer 1 apical tuft dendrites of layer 5B pyramidal tract type neurons. Journal of Neuroscience, 38(41), 8787–8797.

16. Han, X., Guo, S., Ji, N., Li, T., Liu, J., Ye, X., … & Peng, H. (2023). Whole human-brain mapping of single cortical neurons for profiling morphological diversity and stereotypy. Science Advance, 2023.

17. Harris, J. A., Mihalas, S., Hirokawa, K. E., Whitesell, J. D., Choi, H., Bernard, A., … & Zeng, H. (2019). Hierarchical organization of cortical and thalamic connectivity. Nature, 575(7781), 195–202.

18. Kalmbach, B. E., Hodge, R. D., Jorstad, N. L., Owen, S., de Frates, R., Yanny, A. M., … & Ting, J. T. (2021). Signature morpho-electric, transcriptomic, and dendritic properties of human layer 5 neocortical pyramidal neurons. Neuron, 109(18), 2914–2927.

19. Lee, B. R., Budzillo, A., Hadley, K., Miller, J. A., Jarsky, T., Baker, K., … & Berg, J. (2021). Scaled, high fidelity electrophysiological, morphological, and transcriptomic cell characterization. eLife, 10, e65482.

20. Lichtman, J. W., Livet, J., & Sanes, J. R. (2008). A technicolour approach to the connectome. Nature Reviews Neuroscience, 9(6), 417–422.

21. Lipovsek, M., Bardy, C., Cadwell, C. R., Hadley, K., Kobak, D., & Tripathy, S. J. (2021). Patch-seq: Past, present, and future. Journal of Neuroscience, 41(5), 937–946.

22. Liu, L., & Qian, P. (2022). Manifold classification of neuron types from microscopic images. Bioinformatics, 38(21), 4987–4989.

23. Luo, L. (2015). Principles of Neurobiology. Garland Science.

24. Manubens-Gil, L., Zhou, Z., Chen, H., Ramanathan, A., Liu, X., Liu, Y., … & Peng, H. (2023). BigNeuron: a resource to benchmark and predict best-performing algorithms for automated reconstruction of neuronal morphology. Nature Methods, 2023.

25. Mehta, K., Goldin, R. F., & Ascoli, G. A. (2023). Circuit analysis of the Drosophila brain using connectivity-based neuronal classification reveals organization of key communication pathways. Network Neuroscience, 7(1), 269–298.

26. Moffitt, J. R., Bambah-Mukku, D., Eichhorn, S. W., Vaughn, E., Shekhar, K., Perez, J. D., … & Zhuang, X. (2018). Molecular, spatial, and functional single-cell profiling of the hypothalamic preoptic region. Science, 362(6416), eaau5324.

27. Muñoz-Castañeda, R., Zingg, B., Matho, K. S., Chen, X., Wang, Q., Foster, N. N., … & Dong, H.W. (2021). Cellular anatomy of the mouse primary motor cortex. Nature, 598(7879), 159–166.

28. Noble, W. S. (2006). What is a support vector machine?. Nature Biotechnology, 24(12), 1565–1567.

29. Oh, S. W., Harris, J. A., Ng, L., Winslow, B., Cain, N., Mihalas, S., … & Zeng, H. (2014). A mesoscale connectome of the mouse brain. Nature, 508(7495), 207–214.

30. Okigawa, S., Yamaguchi, M., Ito, K. N., Takeuchi, R. F., Morimoto, N., & Osakada, F. (2021). Cell type-and layer-specific convergence in core and shell neurons of the dorsal lateral geniculate nucleus. Journal of Comparative Neurology, 529(8), 2099–2124.

31. Peng, H., Hawrylycz, M., Roskams, J., Hill, S., Spruston, N., Meijering, E., & Ascoli, G. A. (2015). BigNeuron: large-scale 3D neuron reconstruction from optical microscopy images. Neuron, 87(2), 252–256.

32. Peng, H., Xie, P., Liu, L., Kuang, X., Wang, Y., Qu, L., … & Zeng, H. (2021). Morphological diversity of single neurons in molecularly defined cell types. Nature, 598(7879), 174–181.

33. Pierret, T., Lavallée, P., & Deschênes, M. (2000). Parallel streams for the relay of vibrissal information through thalamic barreloids. Journal of Neuroscience, 20(19), 7455–7462.

34. Purves, D., Augustine, G. J., Fitzpatrick, D., Hall, W., LaMantia, A. S., & White, L. (2019). Neurosciences. De Boeck Supérieur.

35. Qu, L., Li, Y., Xie, P., Liu, L., Wang, Y., Wu, J., … & Peng, H. (2022). Cross-modal coherent registration of whole mouse brains. Nature Methods, 19(1), 111–118.

36. Ramón y Cajal, S. (1909) Histologie Du Système Nerveux de L’homme & Des Vertébrés., (Paris: Maloine [Translated by N. Swanson and L.W. Swanson, Oxford University Press, 1995], 1909).

37. Rees CL, Moradi K & Ascoli GA. (2017). Weighing the evidence in Peters’ rule: does neuronal morphology predict connectivity?. Trends in neurosciences, 40(2), 63–71.

38. Russ, D. E., Cross, R. B. P., Li, L., Koch, S. C., Matson, K. J., Yadav, A., … & Levine, A. J. (2021). A harmonized atlas of mouse spinal cord cell types and their spatial organization. Nature Communications, 12(1), 5722.

39. Scala, F., Kobak, D., Bernabucci, M., Bernaerts, Y., Cadwell, C. R., Castro, J. R., … & Tolias, A.S. (2021). Phenotypic variation of transcriptomic cell types in mouse motor cortex. Nature, 598(7879), 144–150.

40. Scheffer, L. K., Xu, C. S., Januszewski, M., Lu, Z., Takemura, S. Y., Hayworth, K. J., … & Plaza, S.M. (2020). A connectome and analysis of the adult Drosophila central brain. eLife, 9, e57443.

41. Scorcioni, R., Polavaram, S., & Ascoli, G. A. (2008). L-Measure: a web-accessible tool for the analysis, comparison and search of digital reconstructions of neuronal morphologies. Nature Protocols, 3(5), 866–876.

42. Sebé-Pedrós, A., Saudemont, B., Chomsky, E., Plessier, F., Mailhé, M. P., Renno, J., … & Marlow, H. (2018). Cnidarian cell type diversity and regulation revealed by whole-organism single-cell RNA-Seq. Cell, 173(6), 1520–1534.

43. Seung, S. (2012). Connectome: How the brain’s wiring makes us who we are. HMH.

44. Sporns, O. (2011). The human connectome: a complex network. Annals of the new York Academy of Sciences, 1224(1), 109–125.

45. Staiger, J. F., & Petersen, C. C. (2021). Neuronal circuits in barrel cortex for whisker sensory perception. Physiological Reviews, 101(1), 353–415.

46. Steinwart, I., & Christmann, A. (2008). Support vector machines. Springer Science & Business Media.

47. Stepanyants, A., Hof, P. R., & Chklovskii, D. B. (2002). Geometry and structural plasticity of synaptic connectivity. Neuron, 34(2), 275–288.

48. Tecuatl, C., Wheeler, D. W., Sutton, N., & Ascoli, G. A. (2021). Comprehensive estimates of potential synaptic connections in local circuits of the rodent hippocampal formation by axonal-dendritic overlap. Journal of Neuroscience, 41(8), 1665–1683.

49. Turner, N. L., Macrina, T., Bae, J. A., Yang, R., Wilson, A. M., Schneider-Mizell, C., … & Seung, H.S. (2022). Reconstruction of neocortex: Organelles, compartments, cells, circuits, and activity. Cell, 185(6), 1082–1100.

50. Van Essen, D. C., Ugurbil, K., Auerbach, E., Barch, D., Behrens, T. E., Bucholz, R., … & WU-Minn HCP Consortium. (2012). The Human Connectome Project: a data acquisition perspective. Neuroimage, 62(4), 2222–2231.

51. Viaene, A. N., Petrof, I., & Sherman, S. M. (2011). Synaptic properties of thalamic input to layers 2/3 and 4 of primary somatosensory and auditory cortices. Journal of Neurophysiology, 105(1), 279–292.

52. Wan, Y., Long, F., Qu, L., Xiao, H., Hawrylycz, M., Myers, E. W., & Peng, H. (2015). BlastNeuron for automated comparison, retrieval and clustering of 3D neuron morphologies. Neuroinformatics, 13, 487–499.

53. Wang, Q., Ding, S. L., Li, Y., Royall, J., Feng, D., Lesnar, P., … & Ng, L. (2020). The Allen mouse brain common coordinate framework: a 3D reference atlas. Cell, 181(4), 936–953.

54. Whitesell, J. D., Liska, A., Coletta, L., Hirokawa, K. E., Bohn, P., Williford, A., … & Harris, J.A. (2021). Regional, layer, and cell-type-specific connectivity of the mouse default mode network. Neuron, 109(3), 545–559.

55. Winnubst, J., Bas, E., Ferreira, T. A., Wu, Z., Economo, M. N., Edson, P., … & Chandrashekar, J. (2019). Reconstruction of 1,000 projection neurons reveals new cell types and organization of long-range connectivity in the mouse brain. Cell, 179(1), 268–281.

56. Yang, J. H., & Kwan, A. C. (2021). Secondary motor cortex: Broadcasting and biasing animal’s decisions through long-range circuits. International Review of Neurobiology, 158, 443–470.

58. Yao, Z., van Velthoven, C. T., Nguyen, T. N., Goldy, J., Sedeno-Cortes, A. E., Baftizadeh, F., … & Zeng, H. (2021). A taxonomy of transcriptomic cell types across the isocortex and hippocampal formation. Cell, 184(12), 3222–3241.

58. Yin, W., Brittain, D., Borseth, J., Scott, M. E., Williams, D., Perkins, J., … & da Costa, N. M. (2020). A petascale automated imaging pipeline for mapping neuronal circuits with high-throughput transmission electron microscopy. Nature Communications, 11(1), 4949.

59. Zeng, H., & Sanes, J. R. (2017). Neuronal cell-type classification: challenges, opportunities and the path forward. Nature Reviews Neuroscience, 18(9), 530–546.

60. Zhang, M., Eichhorn, S. W., Zingg, B., Yao, Z., Cotter, K., Zeng, H., … & Zhuang, X. (2021). Spatially resolved cell atlas of the mouse primary motor cortex by MERFISH. Nature, 598(7879), 137–143.

61. Zhang, M., Pan, X., Jung, W., Halpern, A., Eichhorn, S. W., Lei, Z., … & Zhuang, X. (2023). A molecularly defined and spatially resolved cell atlas of the whole mouse brain. bioRxiv, 2023-03.

62. Zhong, S., Ding, W., Sun, L., Lu, Y., Dong, H., Fan, X., … & Wang, X. (2020). Decoding the development of the human hippocampus. Nature, 577(7791), 531–536.

63. Zingg, B., Hintiryan, H., Gou, L., Song, M. Y., Bay, M., Bienkowski, M. S., … & Dong, H.W. (2014). Neural networks of the mouse neocortex. Cell, 156(5), 1096–1111.

## References (Methods section only)

1. Brown, M. J., Holland, B. R., & Jordan, G. J. (2020). hyperoverlap: Detecting biological overlap in n-dimensional space. Methods in Ecology and Evolution, 11(4), 513–523.

2. Caliński, T., & Harabasz, J. (1974). A dendrite method for cluster analysis. Communications in Statistics-theory and Methods, 3(1), 1–27.

3. Cohen, L., Koffman, N., Meiri, H., Yarom, Y., Lampl, I., & Mizrahi, A. (2013). Time-lapse electrical recordings of single neurons from the mouse neocortex. Proceedings of the National Academy of Sciences, 110(14), 5665–5670.

4. Cortes, C., & Vapnik, V. (1995). Support-vector networks. Machine Learning, 20, 273–297.

5. Delaunay, B. (1934). Sur la sphere vide. Izv. Akad. Nauk SSSR, Otdelenie Matematicheskii i Estestvennyka Nauk, 7(793-800), 1–2.

6. Edelsbrunner, H., & Mücke, E. P. (1994). Three-dimensional alpha shapes. ACM Transactions on Graphics (TOG), 13(1), 43–72.

7. Gong, H., Xu, D., Yuan, J., Li, X., Guo, C., Peng, J., … & Luo, Q. (2016). High-throughput dual-colour precision imaging for brain-wide connectome with cytoarchitectonic landmarks at the cellular level. Nature communications, 7(1), 12142.

8. Han, J., Pei, J., & Tong, H. (2022). Data mining: concepts and techniques. Morgan kaufmann.

9. Iascone, D. M., Li, Y., Sümbül, U., Doron, M., Chen, H., Andreu, V., … & Polleux, F. (2020). Whole-neuron synaptic mapping reveals spatially precise excitatory/inhibitory balance limiting dendritic and somatic spiking. Neuron, 106(4), 566–578.

10. Jiang, S., Guan, Y., Chen, S., Jia, X., Ni, H., Zhang, Y., … & Gong, H. (2020). Anatomically revealed morphological patterns of pyramidal neurons in layer 5 of the motor cortex. Scientific reports, 10(1), 7916.

11. Karlsson, T. E., Smedfors, G., Brodin, A. T., Åberg, E., Mattsson, A., Högbeck, I., … & Olson, L. (2016). NgR1: a tunable sensor regulating memory formation, synaptic, and dendritic plasticity. Cerebral cortex, 26(4), 1804–1817.

12. Lin, H. M., Kuang, J. X., Sun, P., Li, N., Lv, X., & Zhang, Y. H. (2018). Reconstruction of intratelencephalic neurons in the mouse secondary motor cortex reveals the diverse projection patterns of single neurons. Frontiers in neuroanatomy, 12, 86.

13. McLachlan, G. J. (1999). Mahalanobis distance. Resonance, 4(6), 20–26.

14. Morelli, E., Ghiglieri, V., Pendolino, V., Bagetta, V., Pignataro, A., Fejtova, A., … & Calabresi, P. (2014). Environmental enrichment restores CA1 hippocampal LTP and reduces severity of seizures in epileptic mice. Experimental neurology, 261, 320–327.

15. Murase, S., Lantz, C. L., Kim, E., Gupta, N., Higgins, R., Stopfer, M., … & Quinlan, E.M. (2016). Matrix metalloproteinase-9 regulates neuronal circuit development and excitability. Molecular neurobiology, 53, 3477–3493.

16. Nooruddin, F. S., & Turk, G. (2003). Simplification and repair of polygonal models using volumetric techniques. IEEE Transactions on Visualization and Computer Graphics, 9(2), 191–205.

17. Schwarz, G. (1978). Estimating the dimension of a model. The annals of statistics, 461-464.

18. Scrucca, L., Fop, M., Murphy, T. B., & Raftery, A. E. (2016). mclust 5: clustering, classification and density estimation using Gaussian finite mixture models. The R journal, 8(1), 289.

83. Smit-Rigter, L. A., Noorlander, C. W., von Oerthel, L., Chameau, P., Smidt, M. P., & van Hooft, J. A. (2012). Prenatal fluoxetine exposure induces life-long serotonin 5-HT3 receptor-dependent cortical abnormalities and anxiety-like behaviour. Neuropharmacology, 62(2), 865–870.

20. Suter, B. A., & Shepherd, G. M. (2015). Reciprocal interareal connections to corticospinal neurons in mouse M1 and S2. Journal of Neuroscience, 35(7), 2959–2974.

21. Yamashita, T., Vavladeli, A., Pala, A., Galan, K., Crochet, S., Petersen, S. S., & Petersen, C. C. (2018). Diverse long-range axonal projections of excitatory layer 2/3 neurons in mouse barrel cortex. Frontiers in neuroanatomy, 12, 33.

